# CXCL10-driven STAT3 signaling programs pathogenic CD4+ T cell responses and limits antiviral immunity during arthritogenic alphavirus infection

**DOI:** 10.64898/2026.06.08.730944

**Authors:** Sainetra Sridhar, Alexander Lemenze, Aileen Chang, Bobby Brooke Herrera

**Affiliations:** Rutgers Global Health Institute, Rutgers University, New Brunswick, NJ, USA; Department of Medicine, Division of Allergy, Immunology, and Infectious Diseases and Child Health Institute of New Jersey, Rutgers Robert Wood Johnson Medical School, New Brunswick, NJ, USA; Department of Pathology, Immunology, and Laboratory Medicine. Rutgers New Jersey Medical School, Newark, NJ, USA; Department of Medicine, George Washington University, Washington, DC, USA

**Keywords:** O’nyong nyong virus, chikungunya virus, Mayaro virus, CXCL10, CXCR3, T cells, arthritis, immunopathogenesis

## Abstract

Arthritogenic alphaviruses, including o’nyong nyong (ONNV), chikungunya virus (CHIKV), and Mayaro (MAYV) viruses, cause acute and chronic inflammatory joint disease, with disease severity and resolution influenced by host age. The immunological mechanisms governing persistent inflammation remain incompletely defined. In an age-stratified murine model we show that ONNV infection induced elevated *Cxcl10* expression and viral persistence in infected tissues, accompanied by preferential accumulation of CD4+ T cells and limited recruitment of CD8+ T cells. This was associated with skewed T cell differentiation, characterized by increased STAT3 phosphorylation and RORγt expression. Infection with CHIKV and MAYV recapitulated these features, supporting a conserved CXCL10-driven pathogenic program across arthritogenic alphaviruses. Perturbation of this axis reduced CD4+ T cell accumulation, altered T cell differentiation states, and decreased tissue viral burden. Thus, we show that a CXCL10-biased immune response is highly proinflammatory but poorly effective in mediating viral clearance, thus setting the stage for chronic disease.

## Introduction

O’nyong nyong virus (ONNV) is a mosquito-transmitted alphavirus that causes an acute febrile illness characterized by debilitating arthralgia and myalgia. True to its name, “joint breaker,” derived from the Acholi language of northern Uganda, ONNV infection can result in severe joint pain that persists beyond the acute phase of infection [1, 2]. While acute symptoms typically resolve within weeks, a subset of patients develop chronic arthritis lasting months to years, indicating a failure to resolve inflammation. Although the incidence of chronic ONNV disease remains poorly defined, clinical features are thought to parallel those of the closely related chikungunya (CHIKV) or Mayaro (MAYV) viruses, where approximately 30-50% of patients experience prolonged arthritic symptoms [2, 3]. Disease severity and chronicity are further influenced by host age, with both pediatric and elderly populations exhibiting increased morbidity, although the immunological basis for these differences remain unclear [4–7].

Alphaviral infections impose substantial global health burden, and therapeutic options remain limited. Two CHIKV vaccines have recently been approved and demonstrate cross-protection against related alphaviruses, but access remains uneven and no antiviral therapies are currently available [8, 9]. Clinical management is therefore largely supportive, relying on analgesics of immunomodulatory agents with variable efficacy [10]. These constraints underscore the need to define the molecular and immunological drivers of alphaviral pathogenesis to inform targeted therapeutic strategies.

Current models of alphavirus arthritis, largely derived from CHIKV infection in humans and mice, suggest that disease results form a combination of ineffective viral clearance and sustained immune cell infiltration [11]. CD8+ T cell responses are thought to be insufficient for viral control, potentially due to impaired antigen presentation or functional exhaustion [12, 13]. In contrast, CD4+ T cells and monocyte-derived populations have been implicated as key drivers of inflammation, tissue damage, and bone erosion, acting through the secretion of pro-inflammatory mediators and, in the case of monocytes, through osteoclastic differentiation [14–16]. These processes occur within a complex inflammatory milieu that includes type 1 interferons, TNFα, IL-1β, CXCL10, CCL2, which regulate immune cell recruitment and activation and contribute to the persistence of tissue inflammation [15, 17].

Therapeutic targeting of these pathways has yielded mixed outcomes. Modulation of CD4+ T cells activation or trafficking using agents such as abatacept or fingolimod reduces tissue inflammation and histopathology but does not consistently improve viral clearance [18–20]. Cytokine-targeted approaches have similarly produced context-dependent effects, with TNFα blockade exacerbating disease in some settings, whereas CCL2-mediated monocyte recruitment can limit bone pathology [16, 21]. Notably, disruption of CXCL10 signaling in CHIKV infection has been associated with reduced tissue pathology and decreased viral burden, effects that have largely been attributed to altered myeloid cell recruitment [22].

Despite these advances, key aspects of alphaviral immunopathogenesis remain unresolved. Most mechanistic studies have focused on CHIKV infection (3-7 days post infection [dpi]), with limited insights into the drivers of chronic inflammation or viral persistence. In addition, how these immune pathways are regulated across host age, a major determinant of disease severity, remains poorly understood. In this study, we address these gaps using an age-stratified model of ONNV infection to define how inflammatory signals shape T cell responses during alphaviral arthritis. We identify CXCL10 as a central regulator of immune imbalance, linking sustained chemokine expression to preferential accumulation and differentiation of pathogenic CD4+ T cells and limited recruitment of CD8+ T cells. This CXCL10-driven program promotes Th1/Th17 (or Tc1/Tc17) transformation of T cells that is conserved across CHIKV and MAYV infection, revealing a shared mechanism of immunopathogenesis. We further show that the induction of STAT3 phosphorylation underlies these changes. Importantly, perturbation of this axis rebalances T cell responses, reduces tissue inflammation, and enhances viral control. Together, these findings establish a unifying model in which CXCL10 orchestrates age-dependent T cell programming that drives alphaviral disease.

## Results

### ONNV infection results in acute footpad swelling and age-related viral persistence up to 30 dpi

Given our age-informed approach to uncovering alphavirus disease mechanisms, we established a murine model using three age groups: 3-week-old (wo), 6-8-wo and >6-month-old (mo) C57BL/6 mice. Animals were infected with 10^5^ plaque forming units (PFU) of ONNV via subcutaneous footpad injection. We initially assessed overt disease manifestations, including ipsilateral footpad swelling and viral burden in the footpad and draining popliteal lymph node.

All three age groups developed measurable footpad swelling. Peak swelling occurred at 6 dpi in the younger groups, while the older mice exhibited peak swelling at 8 dpi (Fig. 1A-B). To evaluate viral persistence beyond the acute phase, we quantified ONNV RNA in the ipsilateral footpad and draining lymph node at 10 dpi (subacute phase) and 30 dpi (chronic phase). Viral RNA remained detectable in both tissues at both time points, with higher viral copy numbers observed in the footpads of older mice (Fig. 1C-D). These findings indicate an age-associated impairment in viral clearance at the site of infection and are consistent with prior observations in CHIKV models and clinical reports linking age to increased disease severity.

**Figure 1:**
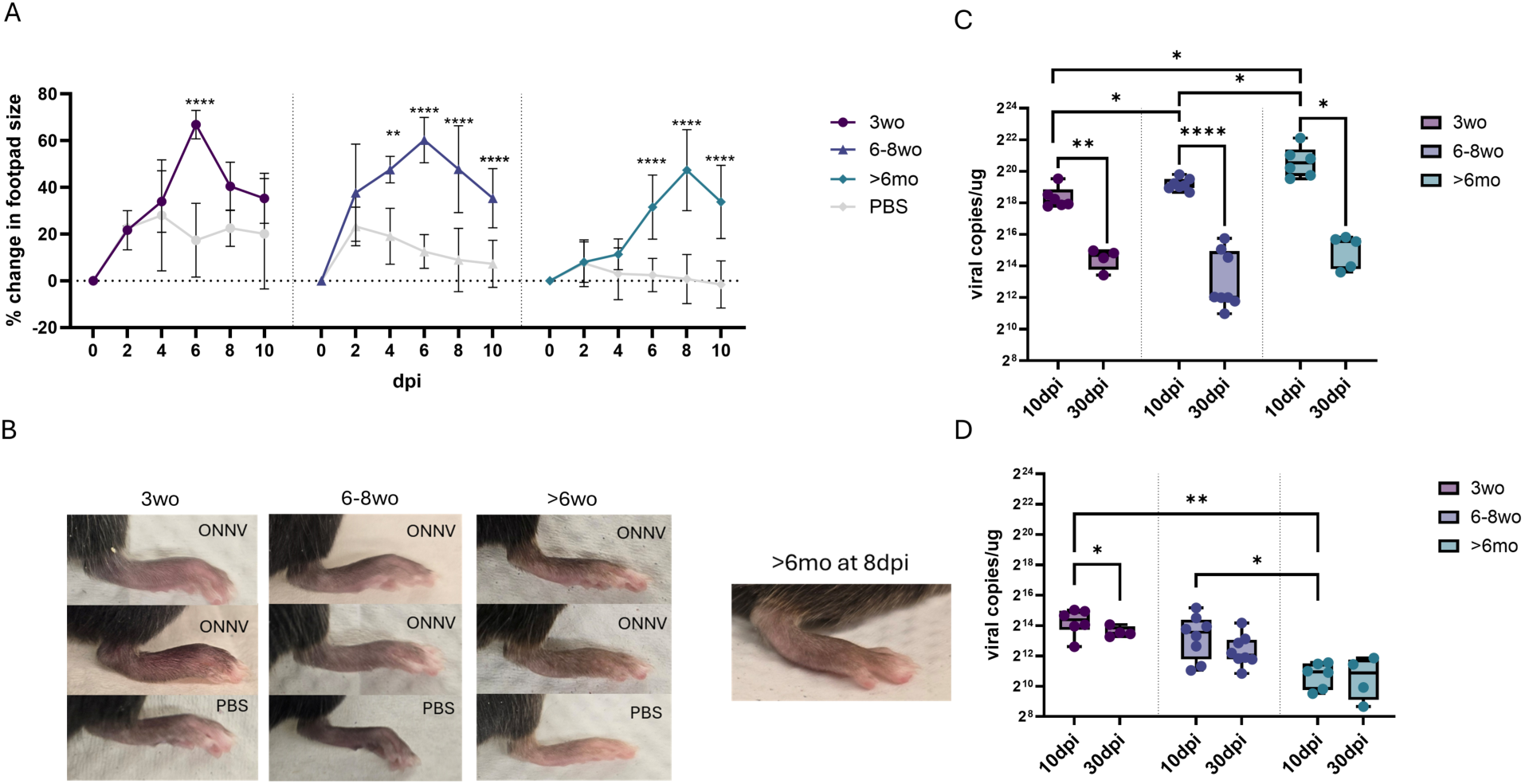
ONNV infection results in acute footpad swelling and age-related viral persistence up to 30 dpi. C57BL/6 mice of the indicated age groups were infected with 10^5 PFU of ONNV or PBS. Footpad swelling over the course of the first 10 dpi was monitored with a digital caliper and percent change in footpad size averaged across N=3-6 mice are plotted (A). Images of the inflamed ipsilateral paws are shown in B. Viral load was measured by qPCR in the ipsilateral footpad (C) and popliteal lymph node (D), where each data point represents an animal (N=4-8). The data were compiled across 3-4 independent experiments. Statistical significance was determined using 2way ANOVA for A and using pairwise T-tests for C and D, ∗*p* < 0.05, ∗∗*p* < 0.01, ∗∗∗*p* < 0.001, ∗∗∗∗*p* < 0.0001.

### ONNV infection is characterized by sustained expression of *Cxcl10* and other pro-inflammatory cytokines and robust CD4+ T cell infiltration in the footpad

Having established a disease phenotype, we next profiled the immune environment in the footpad and draining lymph node at 10 and 30 dpi. We focused on the expression of key pro-inflammatory mediators and the composition of infiltrating immune cells. qRT-PCR analysis of footpad tissue revealed expression of multiple pro-inflammatory cytokines, including *Tnfα*, *Mcp1*, with particularly strong induction of *Cxcl10* at 10 dpi that persisted at 30 dpi, especially in the older mice (Fig. 2A-C). Similar inflammatory signatures were observed in the draining lymph node at 10 dpi (Supplementary Fig. 1A-C).

**Figure 2:**
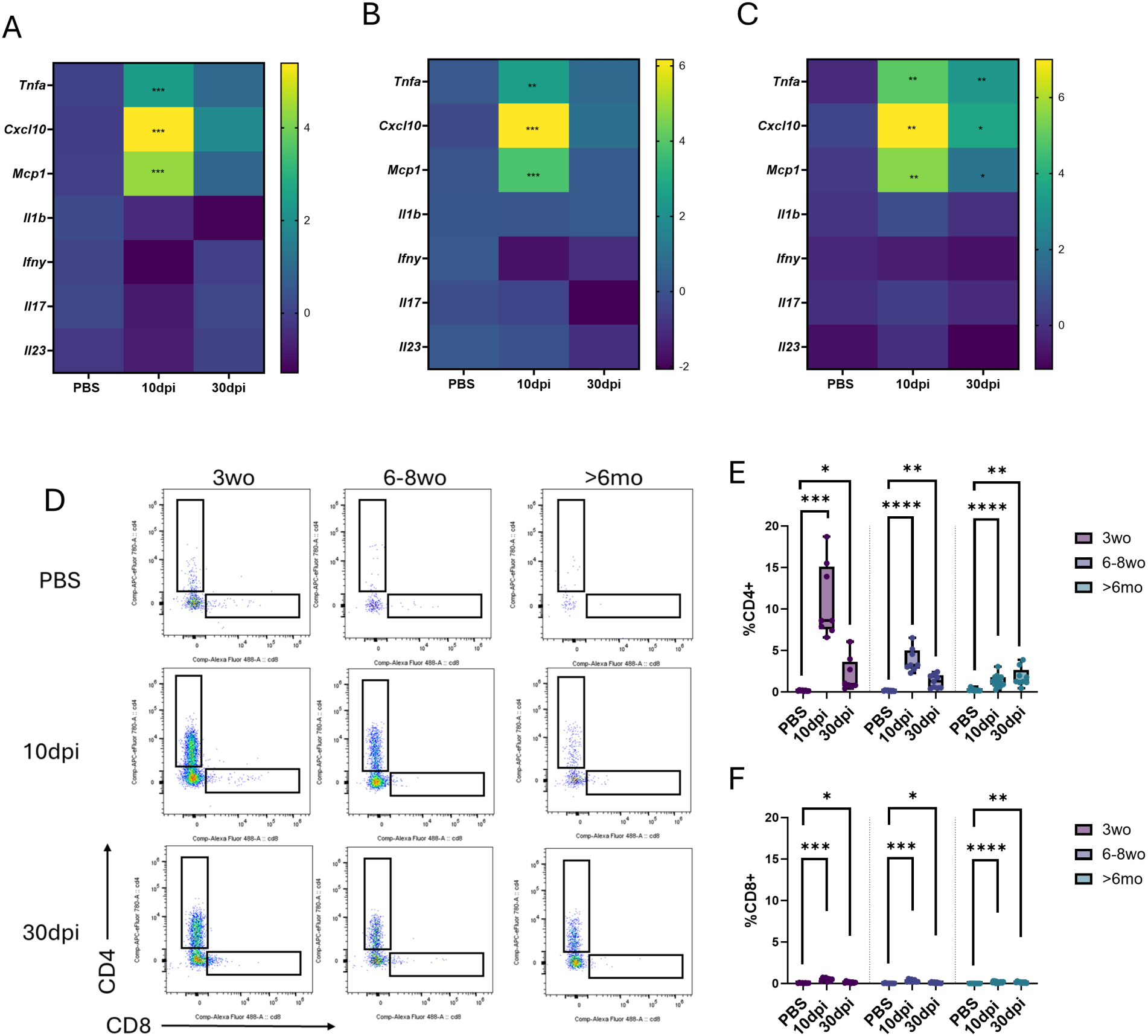
ONNV infection is characterized by sustained expression of *Cxcl10,* robust CD4+ T cell infiltration in the footpad. Mice of the indicated age groups were infected with ONNV or PBS and footpads and lymph nodes were collected at either 10dpi or 30dpi. (A) 3wo, (B) 6-8wo and (C)>6mo mice footpads were subjected to qPCR and the average (across N=5-9) Log2-transformed fold change in expression of various genes relative to uninfected controls is shown. Flow cytometry was used to detect infiltrating CD4+ and CD8+ T cells in the ipsilateral footpads (D-F). D shows representative flow cytometry plots gated on CD3+ T cells, and E and F quantify percentage of CD4+ and CD8+ cells out of total live cells respectively, where each data point represents an animal (N=8-14). Statistical significance was determined using two-sample T-tests where ∗*p* < 0.05, ∗∗*p* < 0.01, ∗∗∗*p* < 0.001, ∗∗∗∗*p* < 0.0001. In A-C, indicated p-values are relative to uninfected controls.

Flow cytometric analysis of footpad infiltrates demonstrated increased frequencies of CD3+ T cells, F4/80+ monocytes, and CD11c+ dendritic cells (DCs) at 10 dpi (Supplementary Fig. 2), consistent with the elevated expression of *Cxcl10* and *Mcp1* in the footpad. Within the T cell compartment, CD4+ T cell proportions were significantly increased in the footpad at 10 dpi, with their continued presence above baseline levels at 30 dpi (Fig. 2D-E). For the older age group, the proportion of CD4+ T cells was higher at 30 dpi compared to 10 dpi (Fig. 2D-E). In contrast, CD8+ T cells, although increased relative to uninfected controls, comprised less than 1% of total cells in the footpad (Fig. 2D, F). The marked enrichment of CD4+ T cells alongside limited CD8+ T cell presence is consistent with prior CHIKV infection suggesting immunopathogenic roles for CD4+ T cells and impaired CD8+ T cells responses [12, 14]. Given that CXCL10 is a potent lymphotactic chemokine, these data suggest that elevated *Cxcl10* may contribute to selective recruitment of T cell subsets to the site of infection [23].

### Fibroblasts and macrophages in the infected footpad produce *Cxcl10* at 10dpi

As an orthogonal method to understand cellular composition at the site of infection, we subjected cells from the ipsilateral footpads of uninfected, 10 dpi and 30 dpi 6-8wo mice to scRNAseq. Using the expression of standard markers for cell types we identified several distinct clusters of cells (Fig. 3A-B). These included footpad associated epithelial cells (clusters 4, 11, 13), fibroblasts (clusters 1, 2, 6, 7, 15), endothelial cells (cluster 5), macrophages (clusters 0, 3, 10, 18), dendritic cells (cluster 14), T cells (cluster 8), B cells (cluster 12), neutrophils (cluster 16), pericytes (cluster 17), and myogenic cells (cluster 9). The elevated presence of myeloid cells (*Cd14* expressing macrophages and DCs) and CD4+ T cells at 10 dpi is in accordance with our flow cytometry observations noted above (Fig. 3B). Furthermore, as noted above, the presence of CD8+ T cells was far limited in comparison to the CD4+ cells and was only revealed upon further sub-clustering the CD3+ cluster (data not shown).

**Figure 3:**
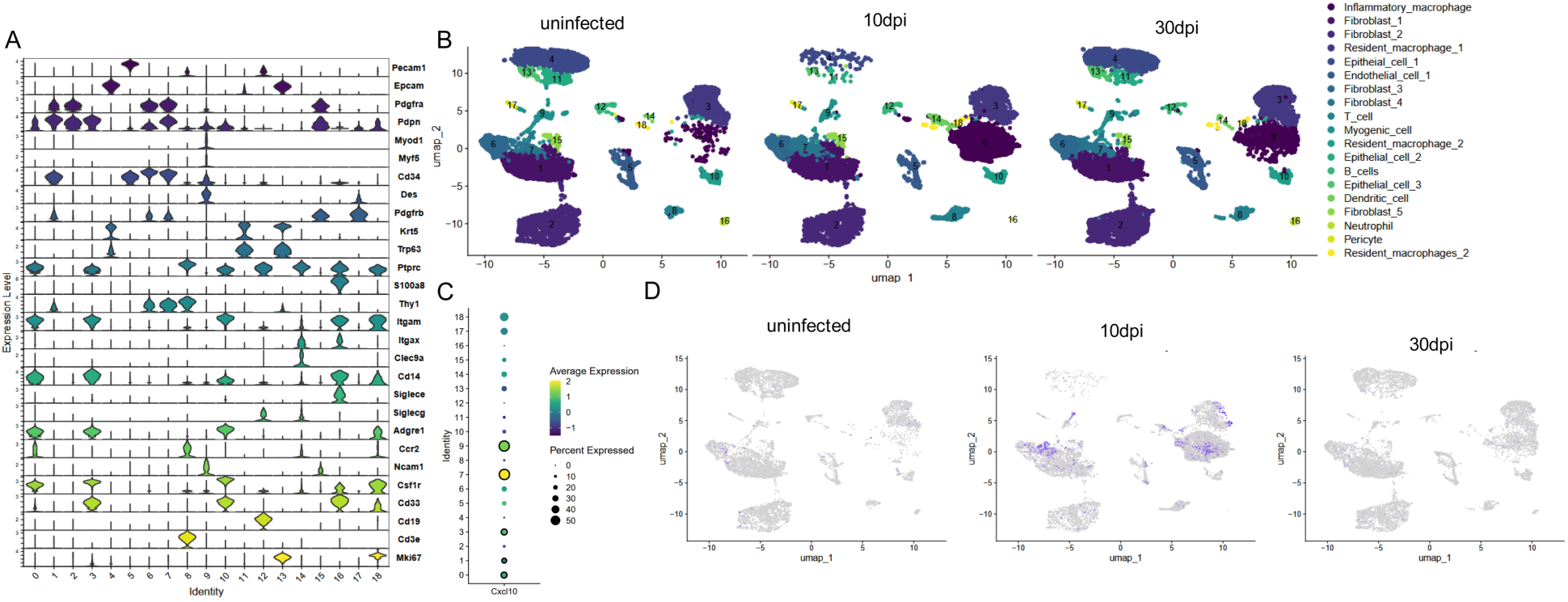
ScRNAseq reveals cxcl10 expression in both resident and infiltrating cells in the footpad at 10dpi. 6-8wo mice were infected with ONNV or with PBS and footpads were collected at 10 and 30 dpi. Footpad cells were fixed and subjected to scRNAseq following the 10x genomics Gem-X flex workflow. Analysis reveals several clusters that were annotated using expression of markers shown in (A). (B) shows a UMAP projection of cell clusters split into uninfected, 10dpi and 30dpi timepoints. C) Measures CXCL10 expression at 10dpi across all 18 clusters. Clusters with statistically significant (P_val_adj<0.05) high fold change increase are indicated with a black border. (D) quantifies the expression levels of CXCL10 across the various clusters and timepoints.

We were interested in identifying the cell types that produce CXCL10 given the above qRT-PCR observation of elevated *Cxcl10* mRNA 10 dpi. The prominent clusters expressing *Cxcl10* at 10 dpi were macrophages (cluster 0,3,18), fibroblasts (6,7), myogenic cells (cluster 9), and pericytes (cluster 17) (Fig. 3C-D). The most significant elevation in expression was in clusters 9, 7, 3 and 0 (Fig. 3C). Cluster 7 corresponds to a Thy1+ fibroblast cluster which are associated with promoting inflammation in rheumatoid arthritis [24]. Among the macrophage clusters, cluster 3 and 18 are tissue resident while cluster 0 corresponds to infiltrating macrophages. It is worth noting that these clusters correspond to cell types known to possess the ability to be infected by alphaviruses and have been speculated to be sources of CXCL10 in CHIKV infections [25–28].

### CD4+ T cells in draining lymph nodes of infected mice exhibit elevated CXCR3 expression

The observed increase in T cell infiltration into the footpad, together with elevated *Cxcl10* expression, prompted us to investigate T cell activation and chemokine receptor expression in the draining lymph node. Using flow cytometry, we first assessed CD44 and CD62L expression to define naïve (CD44-CD62L+), effector (CD44+CD62L-), and central memory (CD44+CD62L+) populations. Both CD4+ and CD8+ T cell compartments exhibited increased proportions of effector T cells following infection, although this expansion was more pronounced in CD4 T cells (Supplementary Fig. 3A-B). To evaluate responsiveness to CXCL10, we quantified expression of CXCR3, the receptor for CXCL10, on total and effector T cell populations. CD4+ T cells showed a significant increase in CXCR3 expression compared to uninfected controls, whereas CD8+ T cells did not exhibit a comparable increase (Fig. 4A and 4C). Among effector populations, CXCR3+ CD4+ T cells comprised approximately 5-10% of the CD4+ compartment at 10 dpi, whereas CXCR3+ CD8+ effector cells remained below 2% (Fig. 4B and 4D). These findings further corroborate a predominantly CD4+ skewed immune response to ONNV infection.

**Figure 4:**
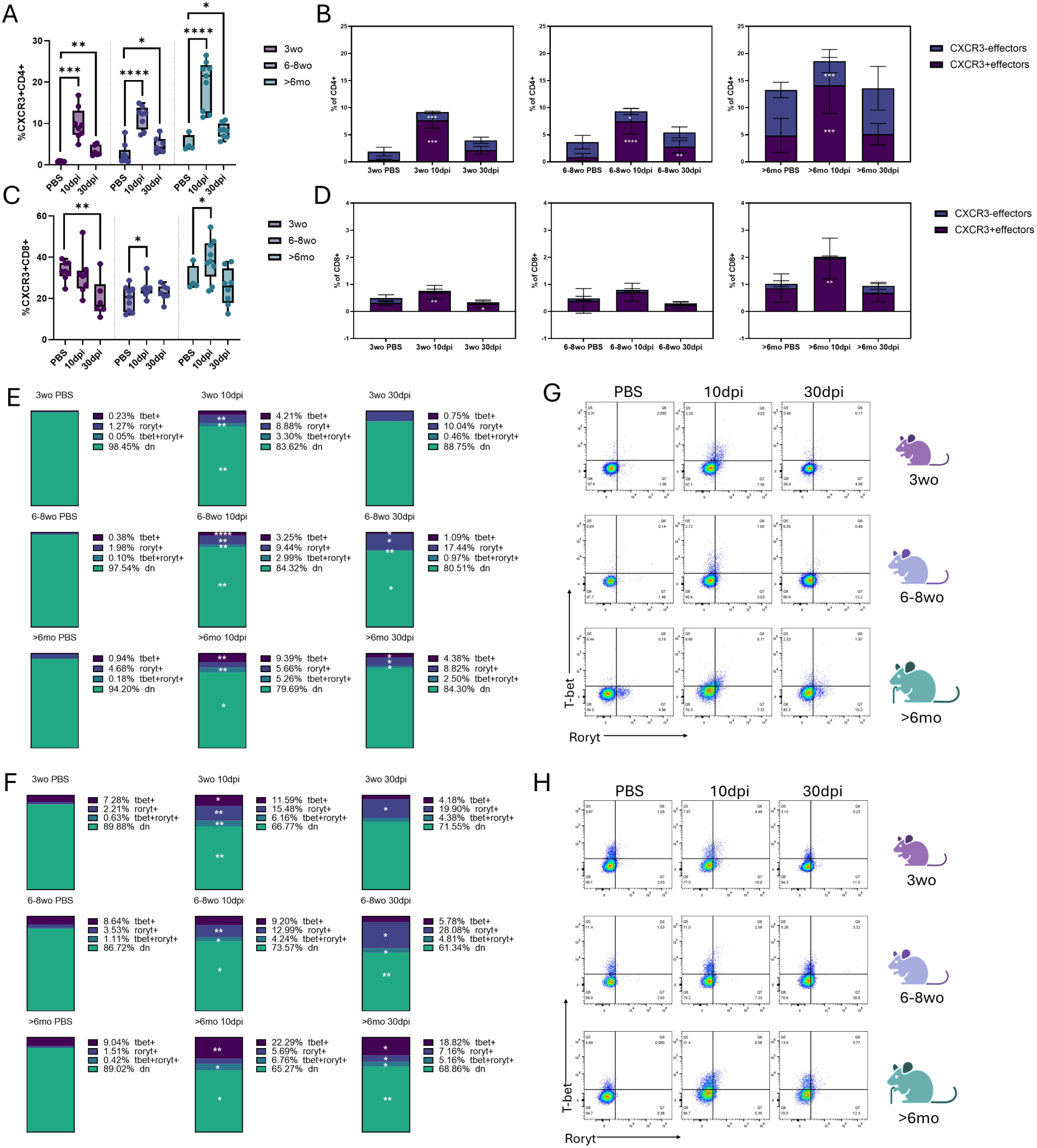
ONNV infection is associated with elevated CXCR3 expression in and pro-inflammatory transcriptional reprogramming of T cells. Mice of the indicated age groups were infected with ONNV or PBS and lymph node cells were collected at 10 or 30dpi and subjected to flow cytometric staining for Tbet and Roryt. (A-D) Percentage of CXCR3+ cells among CD4+ and CD8+ cells in the lymph nodes is shown in A and C respectively (N=7-12). B and D show average percentage (Across N=7-12) of CD4+ and CD8+ cells that are e_ectors (CD44+ CD62L-) stratified by CXCR3 expression. E and F respectively show average frequencies (across N=4-6) of CD4+ or CD8+ T cells that are Tbet+, Roryt+, double positive or double negative. G and H show representative staining for Tbet and Roryt from each group, gated on CD4+ (G) or CD8+ (H) cells. For A-F statistical significance was calculated using Two-sample T tests and the indicated significance is relative to uninfected controls, where ∗*p* < 0.05, ∗∗*p* < 0.01, ∗∗∗*p* < 0.001, ∗∗∗∗*p* < 0.0001.

Interestingly, the highest levels of effector differentiation and CXCR3 expression were observed in older mice, despite their relatively reduced T cell accumulation in the footpad. One possible explanation for this discrepancy is impaired trafficking or functional capacity, potentially due to immune dysfunction or exhaustion [29, 30]. Consistent with this, both CD4+ and CD8+ T cells exhibited increased expression of PD1 and CLTA4 at 10 dpi, with sustained expression at 30 dpi, particularly in older animals (Supplementary Fig. 4). It is also notable that, despite their limited presence in the footpad, a higher proportion of CD8+ T cells expressed CXCR3 compared to CD4+ T cells. This suggest that CXCL10-CXCR3 signaling may differentially influence T cell fate and function, rather than simply recruitment [31].

### Increased *Cxcl10* expression during ONNV infection is associated with pro-inflammatory transcriptional reprogramming of T cells

The CXCL10-CXCR3 signaling axis has been implicated in promoting Th1/Th17 and Tc1/Tc17 differentiation [31–35]. To determine whether elevated CXCL10 and CXCR3 expression was associated with transcriptional reprogramming of T cells in our model, we assessed expression of the lineage-defining transcription factors T-bet and RORγt.

In both CD4+ and CD8+ T cells, we observed an increase in T-bet+ and Rorγt+ populations at 10 dpi, along with a smaller population co-expressing both transcription factors (Fig. 4A, 4C and 4B, 4D respectively). At 30 dpi, the profile shifts towards more RORγt+ cells. Age-dependent differences were also observed, where, at 10 and 30 dpi, older mice exhibited higher frequencies of T-bet+ and T-bet+Rorγt+ T cells compared to younger groups. And in contrast to the younger groups, CD8+ cells of the older mice are more T-bet+ than RORγt+ at both 10 and 30 dpi.

Although cytokine production was not directly measured at the single-cell level in these experiments, elevated expression of *Ifnγ, Tnfα and Il-23* in lymph node tissues at 10 and 30 dpi supports the presence of Th1- and Th-17-like responses (Supplementary Fig. 1). The presence of T-bet+ RORγt+ cells, together with the increased proportion of RORγt+ cells at 30 dpi, suggests phenotypic plasticity between Th1/Tc1 and Th17/Tc17 lineages, accompanied by a shift in effector programming over the course of infection.

### CXCL10-associated T cell reprogramming is conserved across arthritogenic alphaviruses

To determine whether these immune features are specific to ONNV or shared across related alphaviruses, we infected 6-8wo mice with CHIKV and MAYV and performed parallel analyses at 10 dpi. Both infections recapitulated key aspects of the ONNV phenotype, including acute footpad swelling (Supplementary Fig.5), elevated *Cxcl10* expression along with other inflammatory mediators (Supplementary Fig. 5B-C), and increased CD4+ T cell infiltration into the footpad, accompanied by their elevated expression of CXCR3 in the popliteal lymph node, with relative absence of CD8+ T cells (Supplementary Fig. 5D-G). Additionally, both CHIKV and MAYV infection induced T-bet and RORγt expression in CD4+ and CD8+ T cells (Supplementary Fig. 5H-I). These findings suggest that CXCL10-associated T cell reprogramming represents a conserved immunological feature across arthritogenic alphaviruses.

**Figure 5:**
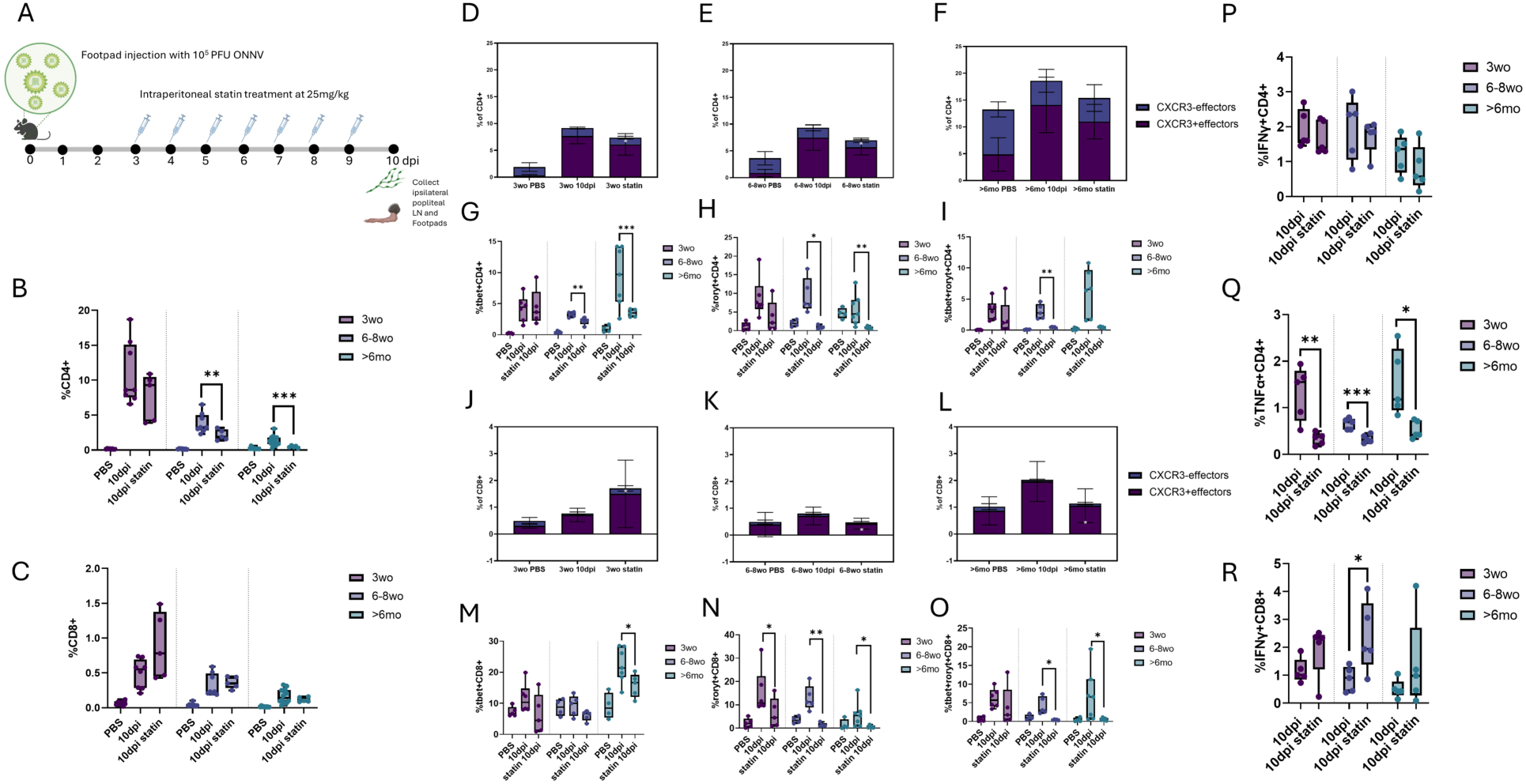
Atorvastatin treatment reduces CD4+ T cell infiltration and attenuates proinflammatory T cell programming during ONNV infection. A is a schematic representation of the atorvastatin treatment strategy after infection of 3wo, 6-8wo or >6mo mice with ONNV. Footpads and lymph nodes were collected for flow cytometric analysis at 10dpi. B and C respectively show frequency of CD4+ and CD8+ cells in the footpad. D-F and J-L quantify average (across N=5-10)frequency of e_ector CD4+ and CD8+ cells in the lymph node, stratified by CXCR3 expression. G-I and M-N show frequency of Tbet+, Roryt+ or double positive CD4+ or CD8+ cells (N=4-7). P-R quantify IFNy+ CD4+, TNFa+CD4+ and IFNy+CD8+ cells in the lymph node after ex vivo stimulation with ONNV-E2-Lfn (N=5-10). Statistical significance was calculated using two-sample T-tests and the plotted significance is relative to uninfected controls unless otherwise indicated, where ∗*p* < 0.05, ∗∗*p* < 0.01, ∗∗∗*p* < 0.001, ∗∗∗∗*p* < 0.0001.

### Inhibition of CXCL10 reduces CD4+ T cell infiltration and attenuates proinflammatory T cell programming

To directly test the role of CXCL10 in driving these immune changes, we inhibited CXCL10 using two complementary approaches: atorvastatin treatment, previously shown to reduce CXCL10 serum levels [36, 37], and administration of a neutralizing anti-CXCL10 antibody. Atorvastatin was administered daily (25 mg/kg) from 3 to 9 dpi, and tissues were analyzed at 10 dpi (Fig. 5A). Anti-CXCL10 treatment consisted of a 200 µg loading dose at 2 dpi followed by 100 µg doses at 4, 6, and 8 dpi (Supplementary Fig. 9A).

Statin treatment across all age groups resulted in reduced infiltration of CD3+ T cells, F4/80+ monocytes, and CD11c+ DCs into the footpad at 10 dpi (Supplementary Fig. 6A). Within the T cell compartment, CD4+ T cell infiltration was significantly reduced relative to untreated controls (Fig. 5B), whereas CD8+ T cell proportions remained largely unchanged (Fig. 5C). In the draining lymph node, statin treatment reduced the frequency of effector (CD44+CD62L-) T cells and decreased CXCR3 expression in both CD4+ and CD8+ populations (Fig. 5D-F, 5J-L). CD4+ T cells also exhibited reduced expression of PD1 and CTLA4 (Supplementary Fig. 7A) and decreased frequencies of T-bet+ and RORγt+ cells (Fig. 5G-I), indicating attenuation of pro-inflammatory differentiation. In contrast, CD8+ T cells primarily showed reduced RORγt expression following statin treatment (Fig. 5L-O), suggesting a shift away from inflammatory programming.

As an added measure of virus-specific T cell function, we performed ex-vivo stimulation of cells isolated from the draining lymph node and quantified cytokine production. For these assays, we used an in-house purified ONNV-specific immunogen consisting of the viral envelope protein E2 fused to a modified, non-infectious domain of the anthrax lethal factor (LFn). Fusion of LFn to immunogenic viral proteins facilitates cytosolic delivery and processing for MHC class I and II presentation, enabling assessment of virus-specific T cell responses without the need for peptide pools [38–43]. Following stimulation, CD4+ T cells from statin-treated mice exhibited reduced frequencies of IFNγ+ and TNFα+ cells compared to untreated controls (Fig. 5P-Q and Supplementary Fig. 8A), consistent with attenuation of their pro-inflammatory phenotype. In contrast, CD8+ T cells from statin-treated mice showed an increase in IFNγ production (Fig. 5R and Supplementary Fig. 8B), suggesting improved effector functionality.

Importantly, treatment with the anti-CXCL10 antibody yielded comparable results to statin treatment, including reduced CD4+ T cell infiltration and attenuation of proinflammatory T cell programming (Supplementary Fig. 9B-O), supporting a direct role for CXCL10 in mediating these effects. Taken together, these data demonstrate that CXCL10-CXCR3 signaling is key driver of T cell immunopathogenesis during alphaviral infection. Perturbation of this axis, either pharmacologically or through direct neutralization, partially reverses CXCL10-associated changes within the T cell compartment, resulting in reduced CD4+ T cell-driven inflammation and a relative shift toward improved CD8+ T cell function.

### CXCL10/CXCR3 drives changes in T-cell differentiation through influencing phosphorylation of STAT3

To better resolve underlying signaling pathways involved in driving CXCL10 associated changes in T cell programming, effector differentiation and antiviral activity, we assessed the presence of pSTAT1/4 and pSTAT3 which are known to drive the differentiation of Th/Tc1 and Th/Tc17 T cells, respectively [32, 33]. Accordingly, we performed phosphoflow cytometry on the lymph node CD4+ and CD8+ T cells from uninfected or 10 dpi mice in all three age groups. Gating on CD4+ T cells, split into T-bet+ and RORγt+ single positive, double positive or double negative cells, we noted the elevation of pSTAT3 mean fluorescence intensity (MFI) in all three age groups and across all sub-groups compared to uninfected controls (Fig. 6B). The increase in pSTAT3 matched the increase in CXCR3 MFI in each category and the highest expression of pSTAT3 was seen in the T-bet+RORγt+ cells (Fig. 6A-B). On the other hand, pSTAT1 and pSTAT4 levels seemed unchanged relative to uninfected controls or sustained a slight increase in the case of the double positive cells (Supplementary Fig. 10 A-B).

**Figure 6:**
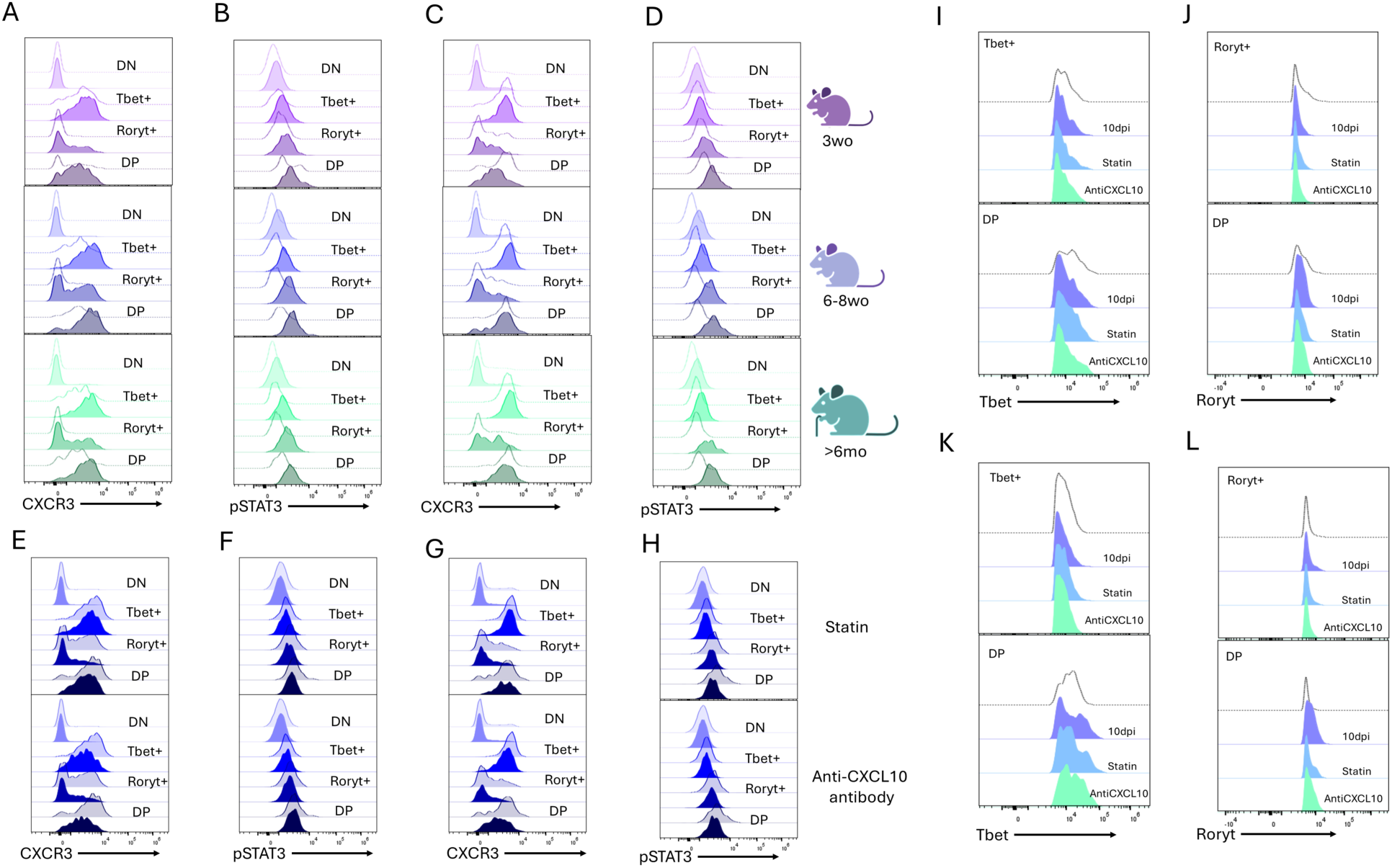
CXCL10/CXCR3 drives changes in T-cell diWerentiation through influencing phosphorylation of STAT3. 3wo, 6-8wo and >6mo mice were infected with ONNV and lymph nodes were collected at 10dpi for flow cytometric analysis. Histograms for the expression of CXCR3 (A, C, E, G), pSTAT3 (B, D, F, H), Tbet (I and K) and Roryt (J and L) were generated by combining expression in cells from N=3 mice. Cells gated on CD4+ (A and B) and CD8+ (C and D) expression and split into Tbet+, Roryt+, double positive (DP) or double negative (DN) categories, and the dotted lines represent corresponding uninfected controls. E-H compares expression in 6-8 wo ONNV infected (translucent colors) and/or either atorvastatin or Anti-cxcl10 antibody treated mice (opaque colors), as indicated, where E-F correspond to CD4+ and G-H correspond to CD8+ cells. I-L compares Tbet and Roryt expression in uninfected (dotted line), ONNV infected (purple), infected and statin treated (blue) or infected and anti-cxcl10 antibody treated (green) CD4+ (I and JJ) and CD8+ (K and L) cells.

The CD8+ T cells followed a similar pattern of expression, with increase in pSTAT3 MFI in all categories (T-bet+, RORγt+, double positive or double negative) at 10dpi compared to their respective uninfected controls in a manner that matched the trend in their respective CXCR3 MFIs (Fig. 6C-D). Interestingly however, the CD8+ cells of 3wo mice showed a smaller relative increase in CXCR3 MFI at 10dpi, compared to the other age groups, accompanied by a smaller induction of pSTAT3 particularly in the T-bet+ single positive subgroup.

This induction of pSTAT3 at 10 dpi was attenuated after atorvastatin or anti-cxcl10 treatment in both CD4+ and CD8+ cells, without much change to pSTAT4 and 1 (Fig. 6E-H and Supplementary Fig. 10E-H). Furthermore, the changes to pSTAT3 seem to accompany changes in T-bet and RORγt MFI within T-bet+, RORγt+ and T-bet+RORγt+ subgroups (Fig. 6I-L).

Specifically, at 10dpi, there is a notable increase in RORγt MFI compared to uninfected controls, in the RORγt+ and double positive populations, that reduces upon statin or anti-CXCL10 antibody treatment (Fig. 6J and 6L). Conversely, the T-bet+ and double positive populations experience a decrease in T-bet MFI at 10 dpi, that increases after atorvastatin or anti-CXCL10 antibody treatment (Fig. 6I and 6K). The former is noted in both CD4+ and CD8+ cells, while the latter is more prominent in CD8+ T cells.

Thus, CXCL10 likely drives changes within the T cell compartment through influencing an increase in pSTAT3, but not pSTAT4 or 1. Despite the previously noted increase in the frequency of T-bet+ cells in both CD4+ and CD8+ cells, the above observations imply that CXCL10/CXCR3 driven increase in STAT3 phosphorylation may strengthen Th17/Tc17 programming at the expense of Th1/Tc1, even in the T-bet+ cells. This further suggests that the CXCL10 dependent increase in pSTAT3 is the likely driving force behind the plasticity between the T-bet and RORγt effector lineages in CD4+ and CD8+ T cells.

### CXCL10 inhibition reduces footpad swelling and viral burden at 10 dpi in an age-related manner

Finally, to determine whether CXCL10-mediated changes in the T cell compartment translated into improved disease outcomes, we evaluated clinical and virological parameters following CXCL10 perturbation. Specifically, we measured footpad swelling and quantified pro-inflammatory cytokine expression and viral RNA levels in the infected footpad at 10 dpi. Statin-treated mice exhibited a reduction in footpad swelling compared to untreated controls, accompanied by decreased expression of multiple pro-inflammatory mediators, including *Cxcl10, Tnfα, and Mcp1* (Fig. 7A-D). Treatment with the anti-CXCL10 antibody produced comparable reductions in footpad swelling and inflammatory cytokine expression; however, as expected, *Cxcl10* transcript levels were not altered due to antibody-mediated neutralization at the protein level (Supplementary Fig. 11 A-B). Consistent with findings in CXCL10-deficient models, attenuation of CXCL10 signaling through either statin treatment or antibody neutralization resulted in a reduction in subacute viral burden within the footpad at 10 dpi (Fig. 7E; Supplementary Fig.11 C) [22]. These data indicate that CXCL10 inhibition not only dampens inflammation but is also associated with improved viral control at the site of infection.

**Figure 7:**
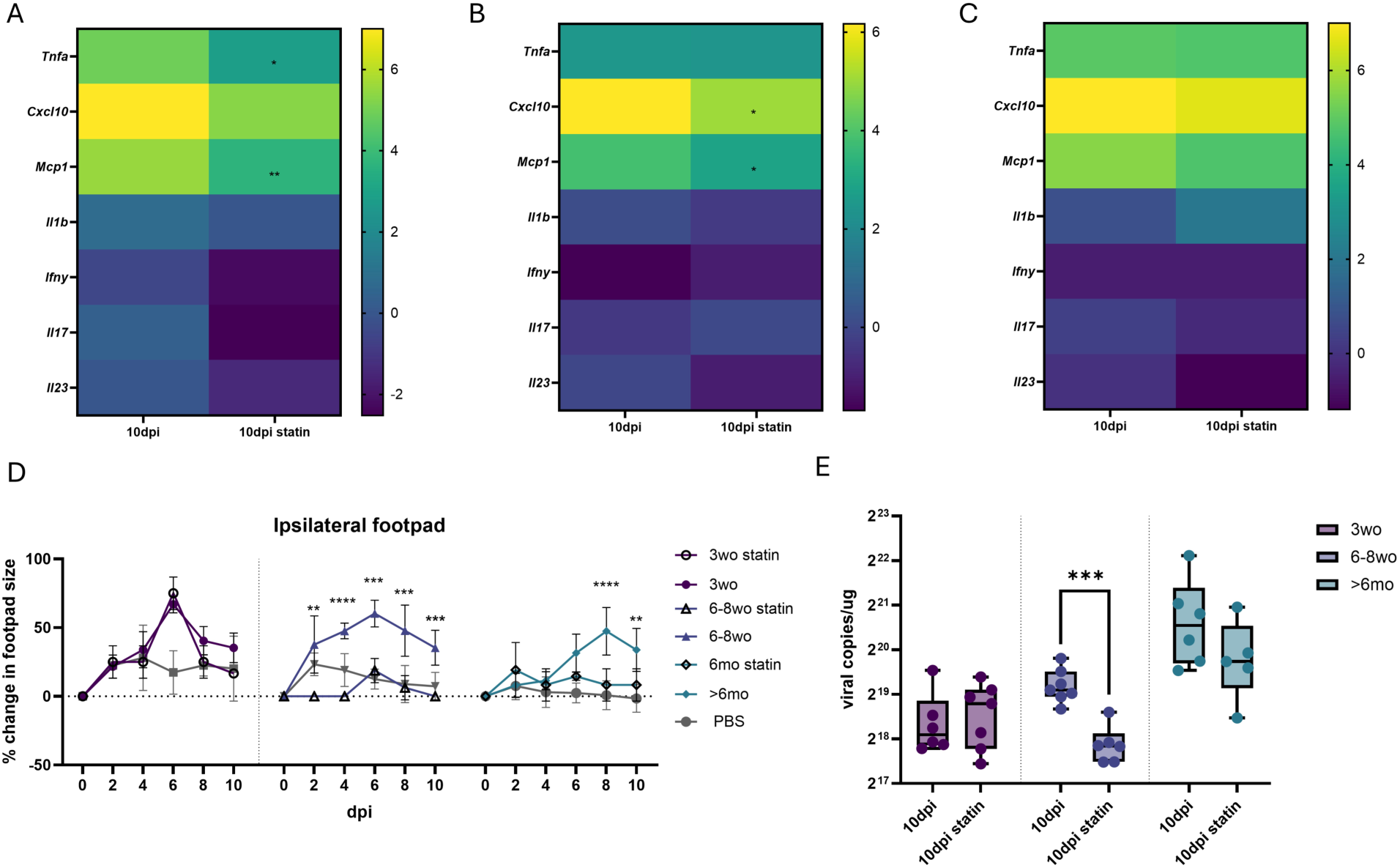
Atorvastatin reduces footpad swelling and viral burden at 10 dpi in an age-related manner. Following infection and atorvastatin treatment, (A) 3wo, (B) 6-8wo and (C)>6mo mice footpads were subjected to qPCR and the average (across N=6-8) Log2-transformed fold change in expression of various genes relative to uninfected controls is shown. D shows change in footpad size following ONNV infection and atorvastatin treatment (N=2-8). E shows footpad viral load measured by qPCR (N=6-8). Statistical significance was calculated using two-sample T-tests and the indicated significance is relative to non-statin treated mice at 10dpi, where ∗*p* < 0.05, ∗∗*p* < 0.01, ∗∗∗*p* < 0.001, ∗∗∗∗*p* < 0.0001.

Notably, these effects were most pronounced in the 6-8-wo and >6-mo age groups, whereas the 3-wo mice exhibited comparatively limited improvement (Fig. 7A and 7E). This differential response suggests that CXCL10-driven immune remodeling operates within an age-dependent immunological context, and that perturbation of this pathway alone is insufficient to fully overcome the broader constraints imposed by the immature immune system in younger hosts. Altogether, these findings support a model in which CXCL10 contributes to both sustained inflammation and impaired viral clearance in an age-dependent manner, linking chemokine-driven immune dysregulation to disease severity in alphaviral infection.

## Discussion

In this study, we define CXCL10 as a central regulator of T cell composition, differentiation, and function during arthritogenic alphavirus infection. Using an age-stratified ONNV model, we demonstrate that elevated *Cxcl10* expression is associated with preferential accumulation and transcriptional programming of CD4+ T cells, limited recruitment and altered functionality of CD8+ T cells, and the persistence of inflammatory responses at sites of infection. These features are conserved across CHIKV and MAYV infection, supporting a unifying model in which CXCL10 orchestrates a pathogenic T cell program that promotes inflammation while constraining effective antiviral immunity.

Our scRNAseq dataset reveals the source of CXCL10 to predominantly be fibroblasts and macrophages in infected footpads. It is speculated that these cells secrete CXCL10 upon being infected with the virus and in a type1 interferon dependent manner [27]. While our dataset does not look directly at whether these cells possessed viral RNA, others have shown that these cell can be infected with alphaviruses [25, 26]. Macrophages in particular have been shown to act as viral reservoirs and play a key role in driving chronic disease [26]. In this study, we were interested in the manner in which they influence the T cell arm of immunity, particularly through their production of CXCL10.

The T cell compartment in our model is characterized by an imbalance between pathogenic and antiviral responses. ONNV infection induced robust infiltration of CD4+ T cells into the infected footpad, whereas CD8+ T cells remained comparatively sparse despite evidence of activation and CXCR3 expression in the draining lymph node. This pattern is consistent with prior work in CHIKV showing that CD4+ T cells contribute to tissue pathology, whereas CD8+ T cell responses are unexpectedly ineffective in clearing virus from infected musculoskeletal sites[44]. Our findings suggest that CXCL10 may be a key component of the mechanism underlying this imbalance. In draining lymph nodes, CD4+ but not CD8+ T cells showed increased CXCR3 expression relative to controls, and the CXCR3+ fraction was particularly enriched within activated and transcriptionally skewed CD4+ subsets. At the same time, CD8+ T cells, despite expressing CXCR3, failed to accumulate efficiently in the footpad and displayed features consistent with impaired function. Taken together, these data support the idea that CXCL10-driven immune organization in alphaviral infection is not simply a matter of chemotactic recruitment but involves differential shaping of CD4+ and CD8+ T cell fates.

The ability of CXCL10 to act as a “driver chemokine” and actively influence T cell activation, polarization, and effector fate through signaling via CXCR3 is well discussed [32]. In several inflammatory settings, CXCL10-CXCR3 signaling has been shown to promote Th1 and Th17 differentiation in CD4+ T cells and analogous Tc1 and Tc17 programs in CD8+ T cells [33–35]. These differentiation states are governed by transcriptional programs centered on T-bet and RORγt and are associated with expression of IFNγ, TNFα, and IL-17 family cytokines [45–47]. Prior work has further linked these programs to CXCR3-associated STAT1/STAT4 and STAT3 activation, which respectively support T-bet- and RORγt-dependent transcriptional states [32, 33]. In our model, we show that CXCL10/CXCR3 seemed to preferentially enhance STAT3 phosphorylation and skew T cells towards a more RORγt controlled fate.

The skew towards RORγt is particularly pertinent given that RORγt+ CD4+ T cells are highly pro-inflammatory and have been implicated in the pathogenesis of autoimmune diseases such as lupus, rheumatoid arthritis, and psoriasis [47, 48]. Th17 cells have also been described in CHIKV patients, where they have been associated with edema, osteoclastogenesis and tissue damage [49–52]. In our mouse model, ONNV infection induced increased frequencies of T-bet+, RORγt+ CD4+ T cells at 10 dpi, including a smaller population co-expressing both transcription factors, consistent with a mixed Th1/Th17 inflammatory state. By 30 dpi, however, the response shifted toward a more RORγt-dominant profile, particularly in the CD4+ compartment, parallelling observations of Th17 enrichment in chronic CHIKV disease [49, 53]. Our data suggests that the CXCL10 driven increase in pSTAT3 and RORγt may set the stage for this transition in the subacute phase of the disease.

At 30 dpi, many of the RORγt+ CD4+ cells were CXCR3 negative, despite persistent inflammation, and this was accompanied by increased *Il-1β* expression in the draining lymph node relative to 10 dpi. One possible interpretation is that early CXCL10-driven T cell activation promotes a mixed Th1/Th17 program during the subacute phase, whereas chronic inflammation becomes increasingly maintained by alternative signals such as IL-1β, which has been implicated in Th17 differentiation [54]. Given the well-established plasticity between Th lineage [55], it is plausible that the RORγt+ cells observed at 30 dpi arise from earlier T-bet+/RORγt+ or Th1/Th17-like intermediates. Defining the temporal relationship between these populations will be important for understanding whether inefficient CXCL10-driven immunity in the subacute phase contributes directly to the emergence of chronic Th17-dominant pathology.

The CD8+ compartment also offers important insight into how CXCL10 may shape antiviral immunity. T-bet expression in CD8+ cells governs short lived effector fates over the formation of memory precursors [56, 57]. STAT3- and RORγt-associated programs in CD8+ T cells, on the other hand, have been linked to impaired cytotoxicity, dysfunctional differentiation, and exhaustion-like states [34, 35]. This framework is relevant to our data, in which CD8+ T cells displayed reduced T-bet and increased RORγt MFIs in lymphoid tissues, were underrepresented in the infected footpad and displayed increased PD1 and CTLA4 expression, particularly in older mice. Importantly, disruption of CXCL10 signaling increased T-bet fluorescence and IFNγ production by CD8+ T cells following antigen-specific stimulation, suggesting that attenuation of this axis improves antiviral effector function. These observations are consistent with findings from chronic LCMV infection, where loss of CXCL10 improved CD8+ T cell functionality and viral control [56], and suggest that CXCL10 may contribute to alphaviral persistence not only by promoting inflammation, but also by biasing CD8+ responses away from effective antiviral activity.

Our perturbation studies support this interpretation. Both atorvastatin and anti-CXCL10 treatment reduced infiltration of CD4+ T cells into the footpad, attenuated pSTAT3 and RORγt expression in the T cell compartment and were associated with reduced tissue inflammation and lower viral burden. While prior studies in CXCL10-deficient CHIKV models attributed improved outcomes primarily to reduced monocyte recruitment [22], our data suggest that reprogramming of T cell responses is also likely to be an important component of this effect. This is particularly relevant because the immunologic changes observed after CXCL10 inhibition were not limited to overall cellularity, but extended to activation state, chemokine receptor expression, transcriptional programming, and virus-specific cytokine responses. In this sense, CXCL10 perturbation does not simply blunt inflammation; it appears to rebalance the immune response away from one dominated by immunopathogenic CD4+ T cells and toward one more permissive of productive CD8+ antiviral function. Importantly, our intention was not to position atorvastatin itself as a specific therapeutic strategy for alphaviral disease per se. Given the pleiotropic effects of statins on metabolism, inflammation, and vascular biology, atorvastatin was used here primarily as an immunologic perturbation approach to interrogate the CXCL10-associated signaling networks underlying T cell dysregulation during infection. The ability of both atorvastatin and direct CXCL10 blockade to partially restore antiviral-associated T cell features while reducing inflammatory pathology supports the broader concept that pathogenic CXCL10-STAT3-RORγt immune programs may be therapeutically targetable. However, future studies will be needed to define more selective interventions capable of modulating these pathways without broadly suppressing protective antiviral immunity.

These mechanistic findings are critically important in the context of the limited treatment options currently available for arthritogenic alphavirus infections. To date, there are no evidence-based treatments for chronic alphaviral arthritis, including arthritis caused by O’nyong-nyong, chikungunya, Mayaro, Ross River, Barmah, and Sindbis viruses, all of which are well-recognized causes of debilitating polyarthritis that can persist for months to years and substantially impair quality of life [58]. Chikungunya arthritis is the most extensively studied of these conditions. For chronic chikungunya arthritis persisting for more than 3 months, empiric therapies such as NSAIDs [59–61], hydroxychloroquine [61–63], sulfasalazine [63], corticosteroids [60], and methotrexate [61–65] have been used. Retrospective studies suggest benefit with methotrexate [61–63, 66], leading to French guidelines recommending its use after 3 months of persistent disease [67]. In clinical practice, initial symptom control often relies on short courses of NSAIDs or low-dose prednisone. However, all current treatment approaches remain empiric, and there is a lack of mechanistically targeted therapies for all arthritogenic alphavirus–associated arthritis. Patients with persistent moderate to severe chronic arthritis are often treated with methotrexate and, in some cases, TNF inhibitors, which carry potential risks including hepatotoxicity, bone marrow suppression, and immunosuppression [64, 68–70]. Statin therapy, and particularly other targeted approaches directed at this newly identified pathway, may offer a more tolerable strategy for modulating disease-relevant mechanisms. Together, these findings support further preclinical and clinical evaluation of CXCL10-pathway modulation as a potential therapeutic strategy across arthritogenic alphavirus infections.

An additional strength of our study is the incorporation of host age as a biological variable. Age is a major determinant of disease severity in alphaviral infection, but mechanistic studies have generally focused on young adult models and acute time points. Here, we show that the effects of ONNV infection and of CXCL10 perturbation are not uniform across age groups. Older mice displayed greater viral persistence, sustained CXCL10 expression, and prolonged maintenance of T-bet-associated T cell states, whereas the youngest mice showed comparatively limited improvement following atorvastatin or anti-CXCL10 antibody treatment. These findings suggest that CXCL10 operates within broader age-dependent immune landscapes rather than as an isolated determinant of pathology. In older hosts, elevated CXCR3 expression and the possibility of higher baseline STAT1/STAT3 activity, as has been described in the setting of inflammaging [71], may reinforce inflammatory circuits and reduce responsiveness to intervention. In contrast, younger hosts may be shaped by a more tolerogenic or immunoregulatory environment [72], potentially limiting both pathogenic T cell differentiation and the therapeutic benefit of suppressing it. The net result is that the same CXCL10-driven pathway may have qualitatively similar but quantitatively distinct consequences across age groups.

At the same time, several limitations warrant consideration. First, our focus on T cell responses does not fully address the contribution of other immune populations, including neutrophils and other granulocytes, which have been implicated in CHIKV pathogenesis and may interact functionally with Th17 cells [49]. Second, while the conserved features observed across ONNV, CHIKV, and MAYV support a shared immune program, the extent to which each virus engages this axis with similar kinetics or tissue specificity remains to be determined. Finally, the mechanisms that sustain chronic inflammation after the subacute CXCL10-rich phase, particularly the potential contribution of IL-1β-driven, CXCR3-negative Th17 states, will require more direct investigation. Our scRNAseq data set, which we mentioned in this text to identify the source of CXCL10, will be invaluable in gaining these insights in future studies.

Overall, our findings place CXCL10 at the center of a broader model of alphaviral immunopathogenesis in which chemokine-driven T cell programming links persistent inflammation to ineffective antiviral immunity. By showing that this program is conserved across arthritogenic alphaviruses and modulated by host age, this work extends the current CHIKV-centered framework and provides a mechanistic basis for heterogeneity in disease outcome. More broadly, these data suggest that successful therapeutic strategies for alphaviral arthritis may need to do more than suppress inflammation; they may need to actively rebalance pathogenic and protective immune programs in a manner informed by host age and disease stage.

## Methods

### Viruses

The three viruses used in this study, O’nyong nyong virus (UgMp30), Chikungunya Virus (181/25) and Mayaro Virus (Guyane) were obtained from BEI resources. Viral stocks were propagated on Vero cells and viral titers were determined using a TCID50 assay, as previously described [40].

### Animal care and in vivo experimental design

All experimental procedures involving animals were conducted in accordance with institutional guidance and approved by the Rutgers, The State University of New Jersey Institutional Animal Care and Use Committee (IACUC) (protocol approval number 202300007). Wild type C57BL/6 mice were purchased from the Jackson Laboratories and colonies were maintained in a pathogen-free facility at the Rutgers Child Health Institute of New Jersey. Since our mouse model is age stratified, we age-matched relevant controls in each age category: 3-week old, 6-8week old and >6month old. No sex based differences were observed and thus results were pooled across male and female animals. The animals did not exhibit any signs of distress upon being infected with the viruses in this study, outside of acute footpad swelling. The experimental endpoints of 10 and 30dpi were selected to represent subacute and chronic timepoints post infection. At these endpoints animals were humanely euthanized using CO2 followed by cervical dislocation before collecting tissues.

*Infection with alphaviruses, monitoring of footpad swelling and treatments with CXCL10 blocking agents:* As part of modelling alphaviral arthritis in vivo, wild type C57BL/6 mice were inoculated via hind footpad subcutaneous injection with 10^5^ Plaque forming units of O’nyong nyong virus (or Chikungunya virus or Mayaro Virus, when relevant). Uninfected controls received an equivalent volume of Phosphate buffered saline (PBS). Ipsilateral footpads of the mice were measured using a digital caliper and photographed at pre-infection (0dpi) and at 2, 4, 6, 8 and 10 dpi.

Atorvastatin (ThermoScientific) and CXCL10 neutralizing antibody (BioXCell) were delivered via intraperitoneal injection. Atorvastatin was delivered at a dose of 25mg/kg daily from 3dpi to 9dpi, similar to previously published studies [22]. For the CXCL10 neutralizing antibody, an initial dose of 200ug was administered at 2dpi, followed by 100ug maintenance dose at 4,6 and 8dpi, similar to previously published studies [73, 74].

At experimental endpoints of 10dpi or 30dpi, animals were humanely euthanized using CO2 followed by cervical dislocation. Ipsilateral popliteal lymph nodes and footpads were collected for further processing for qRT-PCR or Flow cytometry.

### Extracting RNA from tissues and qRTPCR

Collected Ipsilateral Lymph nodes and footpads were stored in RNAlater (ThermoFisher) and held at -80°C until RNA extraction. RNA extraction was performed using the Qiagen RNeasy lipid tissue mini kit following manufacturer’s instructions. Briefly, tissues were homogenized in 1 mL of QIAzol lysis reagent. Lymph nodes were homogenized using a Qiagen Tissue Ruptor II, while footpads were homogenized using a Dounce homogenizer. This was followed by phase separation using chloroform and purification of RNA on spin columns. RNA concentration and purity was determined using a Nanodrop one spectrophotometer (ThermoFisher).

qRTPCR was performed as previously described [39]. In brief, RNA was reverse transcribed using the High-Capacity cDNA Reverse Transcription Kit following manufacturer’s instructions. qPCR to determined viral load and expression of cytokines were conducted using primers listed in Table S1 and Powertrack Sybrgreen QPCR Kits (ThermoFisher). A QuantStudio 6 (ThermoFisher) device was used to acquire qPCR data. Viral copies/ug were determined using a standard curve and fold change in the expression of cytokines was determined using the ΔΔCt method with *Gapdh* serving as the control and then normalizing to relevant uninfected controls.

### Harvesting cells from tissues for flow cytometry and scRNAseq

Extraction of cells from lymph nodes involved passing the tissues through a 70uM mesh filter, washing and pelleting cells in RPMI (supplemented with 10% FBS and 1% antibiotics) at 300x g for 3 minutes. The footpads on the other hand were first digested for 4 hours at 37°C and 5%CO2, shaking at 225 RPM, in 3 mL of RPMI supplemented with 60U of collagenase, 6U of Dispase and 25U DNase, following a protocol adapted from previously published work [20]. Following this, the digested footpads were passed through a 70uM filter, washed and pelleted at 300x g for 3 min and finally passed through a 40uM filter to eliminate any debris. Cells were counted using trypan blue (ThermoFisher) and a Countess III automated cell counter (ThermoFisher).

### scRNAseq sample preparation and analysis

scRNAseq was performed using 10x genomics Gem-X flex expression workflow. 1 million footpad cells were subjected to fixation using the 10x genomics Gem-X flex V2 sample preparation kit, following manufacturer’s instructions. Fixed cells were shipped to Admera Health, New Jersey for library preparation and sequencing.

Raw sequencing reads were aligned to the mouse reference genome (GRCm39) and demultiplexed by cell barcode using Cell Ranger (v10.0.0). Low-quality and outlier cells were removed using the following criteria: >10% mitochondrial reads, >45% ribosomal reads, >2% hemoglobin reads, >5 median absolute deviations (MADs) for the proportion of reads in the top 50 genes, total reads, or detected genes, as well as <100 detected genes or <1000 total reads. Counts were normalized using scTransform, and samples were integrated using reciprocal PCA (RPCA) in Seurat (v5). Cell type annotations were assigned through a combination of SingleR and manual review of canonical markers among the top differentially expressed genes per cluster.

### LFn immunogens and ex vivo stimulation

ONNV envelope protein E2 was cloned into an LFn expression vector, expressed and purified as previously described [39]. For ex vivo stimulation, 200,000-500,000 lymph node cells were plated in RPMI and were treated with purified ONNV-E2-Lfn at 10ug/mL or were left untreated for 16 hours or overnight at 37°C and 5% CO2. After the overnight incubation, Brefeldin A (ThermoFisher) was added to the cells followed by 4 hour incubation, before flow cytometric staining.

### Flow cytometry

Cells from the lymph node or footpad were subjected to flow cytometric staining as previously described [39, 40]. Briefly, this process first involved staining for viability using Live/Dead Red, Blocking of Fc receptors using Normal mouse serum and Anti-CD16/CD34 cocktail (ThermoFisher). Extracellular staining with several fluorophore-conjugated antibodies listed in the Key Resources Table (CD3, CD4, CD8, CD44, CD62L, CXCR3, PD1, F480, CD11c). Cells were then Fixed and Permeabilized using the ebiosciences Foxp3 staining kit (ThermoFisher). Intracellular staining for transcription factors and cytokines was done with several fluorophore-conjugated antibodies listed above (T-bet, RORγt, CTLA4, IFNγ, TNFα, pSTAT3, pSTAT4, pSTAT1). A Cytek Aurora was used to acquire flow cytometric data and analysis was performed using FlowJo software.

### Statistics

In each figure, data were compiled from 3-4 independent experiments. Statistical significance for changes in footpad size (shown in figures 1A and 7D, Supplementary figures 5A and 11C) were calculated using a 2way ANOVA test, run on Prism. For fold changes in RNA expression determined through qPCR and the flow cytometry data, two-sample heteroscedastic one-tailed T-tests were conducted between uninfected and 10dpi or 30dpi samples or between 10dpi and statin or anti-cxcl10 antibody treated samples, as indicated in the figure captions. For the scRNAseq derived differences in *Cxcl10* expression in various clusters, fold changes and p-values were determined using the FindMarkers function in Seurat, comparing uninfected and 10dpi conditions in each cluster.

## Resource Availability

### Lead contact

Further information and requests for resources and reagents should be directed to and will be fulfilled by the lead contact, Bobby Brooke Herrera.

### Materials availability

This study did not generate new unique reagents.

### Data and code availability

- All data supporting the findings of this study are available within the article and its supplemental information. Raw flow cytometry files and RT-qPCR results are available from the lead contact upon reasonable request.
- This study did not generate any original code. scRNAseq data have been deposited at GEO and the accession number is available in the Key Resources table.
- No additional resources were generated or analyzed during this study.

## Acknowledgments

We would like to thank Rutgers Global Health Institute, Rutgers Robert Wood Johnson Medical School, and the Child Health Institute of New Jersey for their support.

## Author contributions

Conceptualization, S.S. and B.B.H.; formal analysis, S.S., A.L., B.B.H.; investigation, S.S., A.L., B.B.H.; resources, B.B.H.; data curation, S.S.; writing – original draft preparation, S.S. and B.B.H.; writing – review and editing, S.S., A.L., B.B.H.; visualization, S.S. and B.B.H.; supervision, B.B.H.; funding acquisition, B.B.H. All authors have read and agreed to the published version of the manuscript.

## Supplemental Information

See document S1: Figures S1-S11 and Table S1.

## STAR★METHODS

### KEY RESOURCES TABLE

| REAGENT OR RESOURCE | SOURCE | IDENTIFIER |
| --- | --- | --- |
| **Antibodies** |  |  |
| CD16/CD32 Fcy III/II (Thermo fisher sciences) | Invitrogen | Catalog # 14-0161-82 |
| CD3 Monoclonal Antibody (OKT3), Alexa Fluor™ 700, eBioscience | Invitrogen | Catalog # 56-0037-42, RRID: AB_10714978 |
| CD4 Monoclonal Antibody (GK1.5), APC-eFluor™<br>780 | Invitrogen | Catalog # 47-0041-80, RRID:<br>AB_11219883 |
| CD8 Monoclonal Antibody (53-6.7) Alexa Fluor 488 | Invitrogen | Cat# 53-008180 |
| CD44 Monoclonal Antibody (IM7), Brilliant Violet™<br>421, eBioscience™ | Invitrogen | Catalog # 404-0441-82,<br>RRID:<br>AB_2925505 |
| CD62L (L-Selectin) Monoclonal Antibody (MEL-14),<br>PE-Cyanine 7 | Invitrogen | Catalog # A14721,<br>RRID:<br>AB_2534237 |
| CD279 (PD-1) Monoclonal Antibody (J43), Super<br>Bright™ 780, eBioscience™ | Invitrogen | Catalog # 78-9985-82,<br>RRID:<br>AB_2734947 |
| CD152 (CTLA-4) Monoclonal Antibody (UC10-4B9), APC, eBioscience™ | Invitrogen | Catalog # 17-1522-82,<br>RRID:<br>AB_2016700 |
| CD183 (CXCR3) Monoclonal Antibody (CXCR3-173), Super Bright™ 600, eBioscience™ | Invitrogen | Catalog # 63-1831-82,<br>RRID:<br>AB_2734898 |
| T-bet Monoclonal Antibody (eBio4B10 (4B10)), PE-Cyanine5, eBioscience™ | Invitrogen | Catalog # 15-5825-82,<br>RRID:<br>AB_2815071 |
| ROR gamma (t) Monoclonal Antibody (AFKJS-9), PE, eBioscience | Invitrogen | Catalog # 12-6988-82,<br>RRID:<br>AB_1834470 |
| ROR gamma (t) Monoclonal Antibody (B2D), PE-Cyanine7, eBioscience™ | AB_2784671 | AB_2784671 |
| Rat anti-mouse IFN gamma Monoclonal Antibody (XMG1.2), eFluor™ 660 | Invitrogen | Catalog# 50-7311-80,<br>RRID:<br>AB_11218909 |
| TNF alpha Monoclonal Antibody (MP6-XT22),<br>Brilliant Violet™ 421, eBioscience™ | Invitrogen | Catalog # 404-7321-80,<br>RRID:<br>AB_2925545 |
| Phospho-Stat1 (Tyr701) Recombinant Rabbit<br>Monoclonal Antibody (Stat1Y701-3E6), APC | Invitrogen | Catalog # MA5-37041,<br>RRID:<br>AB_2896976 |
| Phospho-STAT3 (Tyr705) Recombinant Mouse<br>Monoclonal Antibody (LUVNKLA), Alexa Fluor™<br>Plus 555 | Invitrogen | Catalog #<br>740067MP555 |
| Phospho-STAT4 (Tyr693) Monoclonal Antibody<br>(4LURPIE), APC, eBioscience™ | Invitrogen | Catalog # 17-9044-42 |
| *InVivo*MAb anti-mouse CXCL10 (IP-10) | Bio X Cell | Catalog #BE0440,<br>RRID:<br>AB_3696163 |
| **Bacterial and virus strains** |  |  |
| NIH: O'nyong nyong Virus (UgMP30) | Human isolate, BEI<br>Resources | Catalog No. NR-51661 |
| NIH: Chikungunya Virus (181/25) | Human isolate, BEI<br>Resources | Catalog No. NR-56523 |
| NIH: Mayaro Virus (Guyane) | Human isolate, BEI Resources | Catalog No. NR-49911 |
| **Biological samples** |  |  |
| **N/A** |  |  |
| **Chemicals, peptides, and recombinant proteins** |  |  |
| LFn-ONNV-E2 | Mir Biosciences, Inc | N/A |
| Brefeldin A Solution | Thermo Fisher Scientific | Cat# 00-4506-51 |
| RBC Lysis Buffer | Thermo Fisher Scientific | Cat# J62150.AP |
| eBioscience™ Foxp3 / Transcription Factor Staining Buffer Set | Thermo Fisher Scientific | Cat# 00-5523-00 |
| Dispase | Gibco | Catalog # 17105041 |
| Collagenase, Type IV | Gibco | Catalog # 17104019 |
| DNase I Solution (2500 U/mL) | Thermo Scientific | Catalog # 90083 |
| Atorvastatin Calcium Trihydrate | Thermo Scientific Chemicals | Catalog # 464810010 |
| UltraComp eBeads™ Plus Compensation Beads | Invitrogen | Catalog # 01-3333-41 |
| **Critical commercial assays** |  |  |
| LIVE/DEAD™ Fixable Red Dead Cell Stain Kit | Thermo Fisher Scientific | Catalog # L34972 |
| RNeasy Lipid Tissue Mini Kit | Qiagen | Catalog # 74804 |
| High-Capacity cDNA Reverse Transcription Kit | Thermo Fisher Scientific | Catalog # 4368814 |
| PowerTrack SYBR Green Master Mix | Applied Biosystems | Catalog # A46109 |
| Gem-X Flex Sample Preparation v2 Kit | 10x Genomics | PN-1000781 |
| **Deposited data** |  |  |
| Mouse footpad scRNAseq | This study | GSE334557 |
| **Experimental models: Cell lines** |  |  |
| Vero cells | ATCC | Catalog # CCL-81.4,<br>RRID: CVCL_0059 |
| **Experimental models: Organisms/strains** |  |  |
| Mice C57BL/6 | The Jackson Laboratory | RRID: MGI:2159769 |
| **Oligonucleotides** |  |  |
| N/A |  |  |
| **Recombinant DNA** |  |  |
| N/A |  |  |
| **Software and algorithms** |  |  |
| Cytek Aurora Spectral Analyzer | Cytek | RRID:SCR_01982<br>6 |
| Cytek SpectroFlo | Cytek | RRID:SCR_02549<br>4 |
| FlowJo version 10.2 | TreeStar, Inc | RRID:SCR_00852<br>0 |
| GraphPad Prism | GraphPad software | RRID:SCR_00279<br>8 |
| QuantStudio Real-Time PCR Software | Applied Biosystems | SCR_021096 |
| BioRender | BioRender | RRID:SCR_01836<br>1 |
| R | R core Team |  |
| Seurat V5.4.0 | Seurat | https://satijalab.org/seurat/<br>RRID:<br>SCR_016341 |
| **Other** |  |  |
| N/A |  |  |

