## Supplementary Information for "CXCL10-driven STAT3 signaling programs pathogenic CD4+ T cell responses and limits antiviral immunity during arthritogenic alphavirus infection"

### Supplementary Tables

**Supplementary Table 1: List of primers used in the qPCR assays described in the text.**

| *Gene* | *Forward primer* | *Reverse primer* | *Reference* |
|---|---|---|---|
| *Onnv* | 5’-CGCAGCTTACGGGTTTCATA-3’ | 5’-GCAACGCCTTCAGAAACGC-3’ | [75] |
| *Tnfa* | 5’-ATGAGAAGTTCCCAAATGGC-3’ | 5’-CTCCACTTGGTGGTTTGCTA-3 | [76] |
| *Cxcl10* | 5’-GCCGTCATTTTCTGCCTCAT-3’ | 5’-GCTTCCCTATGGCCCTCATT-3’ | [76] |
| *Mcp1* | 5'-GCATCCACGTGTTGGCTCA-3' | 5'-CTCCAGCCTACTCATTGGGATCA-3' | [76] |
| *Il1b* | 5’-CACAGCAGCACATCAACAAG-3’ | 5’-GTGCTCATGTCCTCATCCTG-3’ | [76] |
| *Ifny* | 5'-AGAGGATGGTTTGCATCTGGGTCA-3' | 5'-ACAACGCTATGCAGCTTGTTCGTG-3' | [78] |
| *il17* | 5’-TCTCCACCGCAATGAAGACC-3’ | 5’-CACACCCACCAGCATCTTCT-3’ | [76] |
| *Il-23p19* | 5'-CCAGCAGCTCTCTCGGAATC-3' | 5'-TCATATGTCCCGCTGGTGC-3' | [77] |

### Supplementary figures

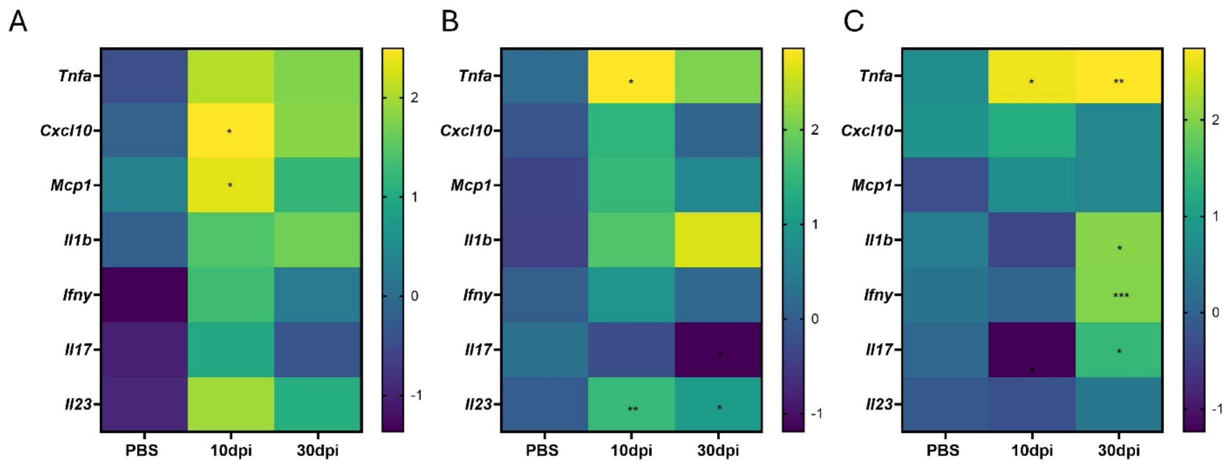


**Supplementary Figure 1: ONNV infection is characterized by sustained expression pro-inflammatory cytokines in the popliteal Lymph node.** Mice of the indicated age groups were infected with ONNV or PBS and lymph nodes were collected at either 10dpi or 30dpi. (A) 3wo, (B) 6-8wo and (C)>6mo mice ipsilateral popliteal lymph nodes were subjected to qPCR and the average (across N=5-9) Log2-transformed fold change in expression of various genes relative to uninfected controls is shown. Statistical significance was determined using two-sample T-tests where $*p < 0.05$, $**p < 0.01$, $***p < 0.001$, $****p < 0.0001$. In A-C, p-values relative to uninfected controls are shown.

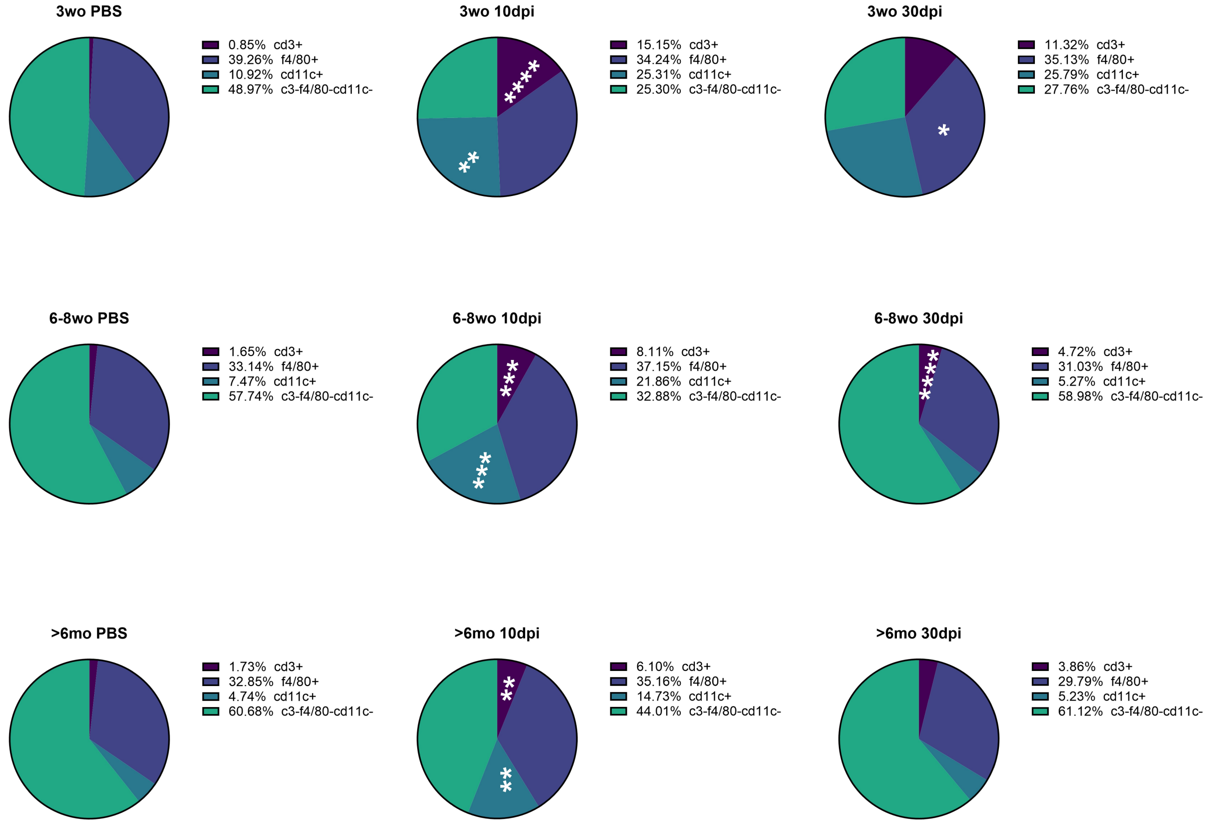


**Supplementary Figure 2: ONNV infection causes influx of T cells, Macrophages and Dendritic cells into the footpad.** Mice of the indicated age groups were infected with ONNV or PBS and footpads were collected at either 10dpi or 30dpi and cells obtained were subjected to flow cytometric analysis. Average frequency (across N=3-8) of CD3+, F480+ (Macrophages) and CD11c+ (Dendritic cells) is plotted above. Two-sample T-tests were used to determining statistical significance and p values shown above are relative to the uninfected mice in each age group, $*p < 0.05$, $**p < 0.01$, $***p < 0.001$, $****p < 0.0001$.

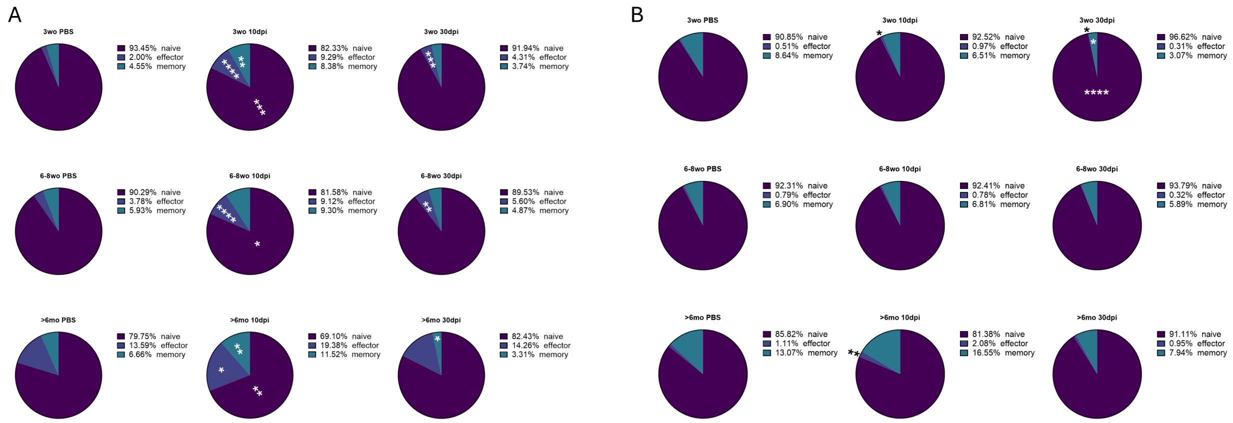


**Supplementary Figure 3: ONNV infection results in increased CD4+ effector cells in the popliteal lymph node, with only limited change to the percentage of CD8+ effectors.** Mice of the indicated age groups were infected with ONNV or PBS and lymph nodes were collected at either 10dpi or 30dpi for flowcytometric analysis. Average frequency (across N=3-8) of naïve (CD44-CD62L+), effector (CD44+ CD62L-) and central memory (CD44+C62L+) CD4+ (A) and CD8+ (B) T cells is shown above. Two-sample T-tests were used to determining statistical significance and p values shown above are relative to the uninfected mice in each age group, $*p < 0.05$, $**p < 0.01$, $***p < 0.001$, $****p < 0.0001$.

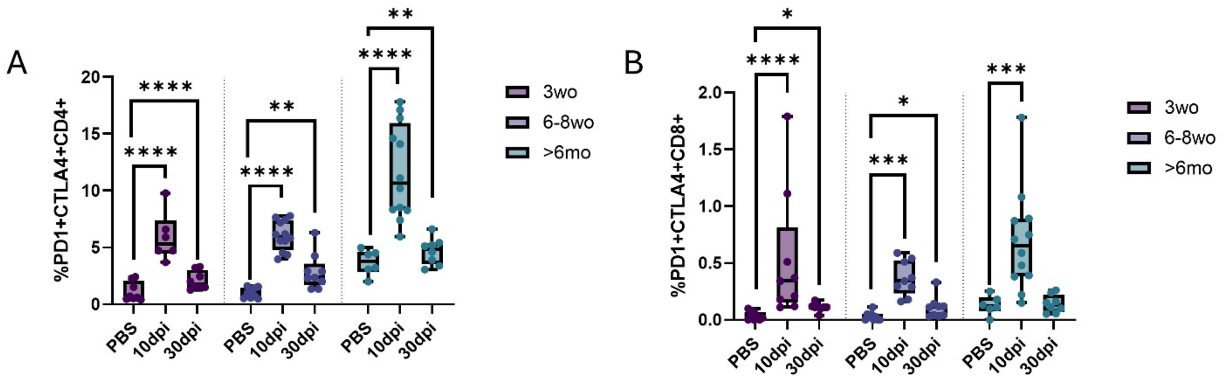


**Supplementary Figure 4: At 10 and 30dpi following ONNV infection, CD4+ and CD8+ T cells express increased PD1 and CTLA4.** Mice of the indicated age groups were infected with ONNV or PBS and ipsilateral lymph nodes were collected at either 10dpi or 30dpi. Flow cytometric staining for PD1 and CTLA4 was done and quantified CD4+PD1+CTLA4+ (A) and CD8+PD1+CTLA4+ (B) T cells are shown above (N=6-12). Two-sample T-tests were used to determining statistical significance and p values shown above are relative to the uninfected mice in each age group, $*p < 0.05$, $**p < 0.01$, $***p < 0.001$, $****p < 0.0001$.

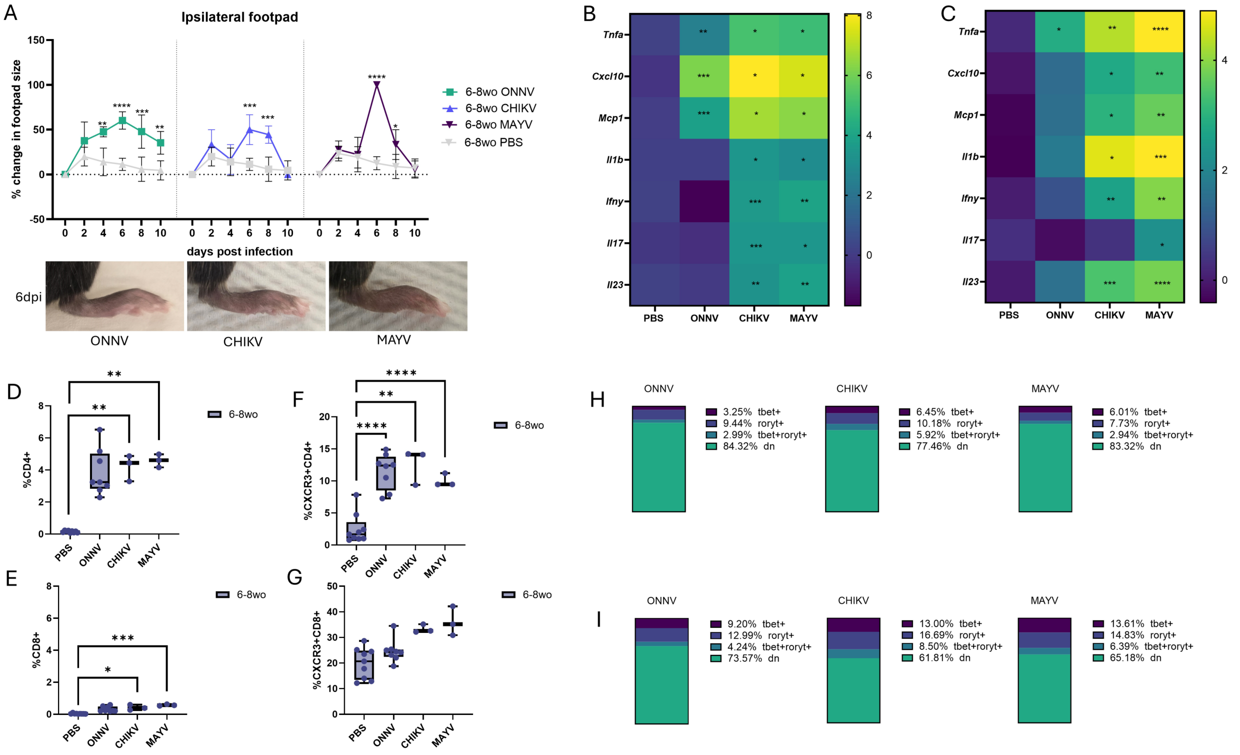


**Supplementary Figure 5: Proinflammatory T cell reprogramming is conserved across arthritogenic alphaviruses.** 6-8wo mice were infected with 10^5 PFU of ONNV, CHIKV or MAYV. Footpad swelling over the course of the first 10 dpi was monitored with a digital caliper and percent change in footpad size averaged across N=3-6 mice are plotted in A along with images of the inflamed paws at 6dpi. Ipsilateral footpads (B) and popliteal nodes (C) were collected at 10dpi and subjected to qPCR and the average (across N=3-7) Log2-transformed fold change in expression of various genes relative to uninfected controls is shown. Flow cytometry was used to measure infiltrating CD4+ and CD8+ T cells in the ipsilateral footpads (D and E). In the lymph nodes, CXCR3 expression (F and G) and Tbet and Roryt expression (H and I) was measured on both CD4+ and CD8+ T cells. In A, statistical significance was calculated using a 2-way ANOVA. In B-I, statistical significance was calculated using two-sample T-tests and the indicated significance is relative to uninfected controls, where $*p < 0.05$, $**p < 0.01$, $***p < 0.001$, $****p < 0.0001$.

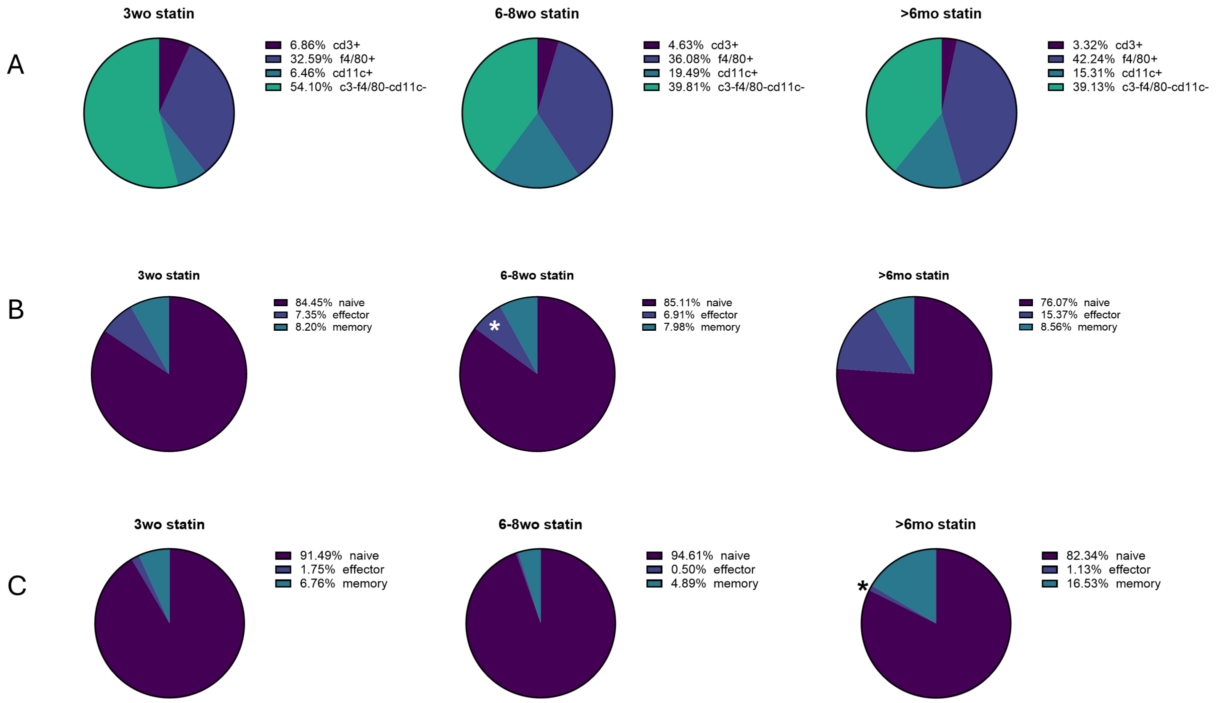


**Supplementary Figure 6: Atorvastatin treatment reduces immune cell T cell infiltration and attenuates proinflammatory T cell programming during ONNV infection.** Footpads and popliteal lymph nodes of ONNV infected and atorvastatin treated mice of the indicated age groups were subjected to flow cytometric analysis. A shows the average frequency (N=5) of CD3+, F480+ (Macrophages) and CD11c+ (Dendritic cells) in the footpads of statin treated mice. B and C show the average frequency (N=5) of naïve (CD44-CD62L+), effector (CD44+ CD62L-) and central memory (CD44+C62L+) CD4+ (B) and CD8+ (C) T cells. Two-sample T-tests were used to determining statistical significance and p values shown above are relative to non-statin treated mice at 10dpi in each age group, $*p < 0.05$, $**p < 0.01$, $***p < 0.001$, $****p < 0.0001$.

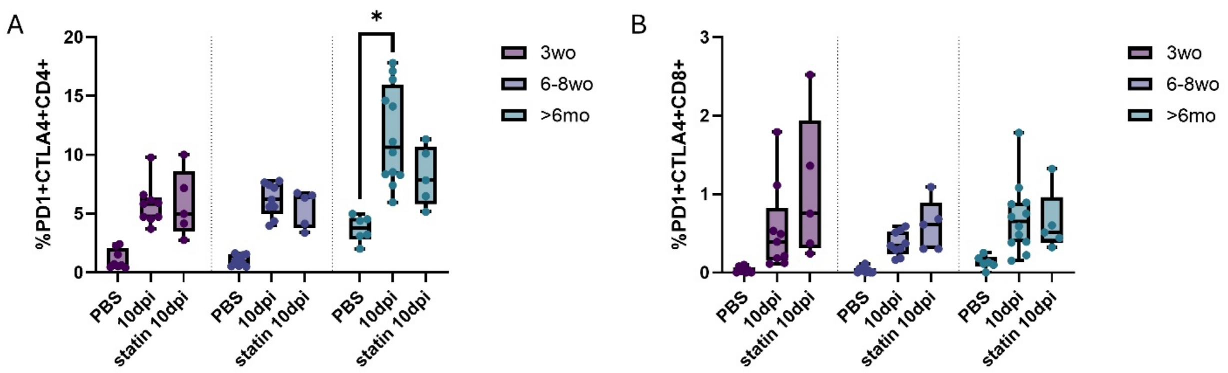


**Supplementary Figure 7: Atorvastatin treatment attenuates PD1 and CTLA4 expression in CD4+ and CD8+ cells.** Mice of the indicated age groups were infected with ONNV and treated with atorvastatin. Lymph node T cells were subjected to flow cytometric staining for PD1 and CTLA4. CD4+PD1+CTLA4+ (A) and CD8+PD1+CTLA4+ (B) T cells are shown above (N=5-10). Two sample T-tests were used to determining statistical significance and $*p < 0.05$, $**p < 0.01$, $***p < 0.001$, $****p < 0.0001$.

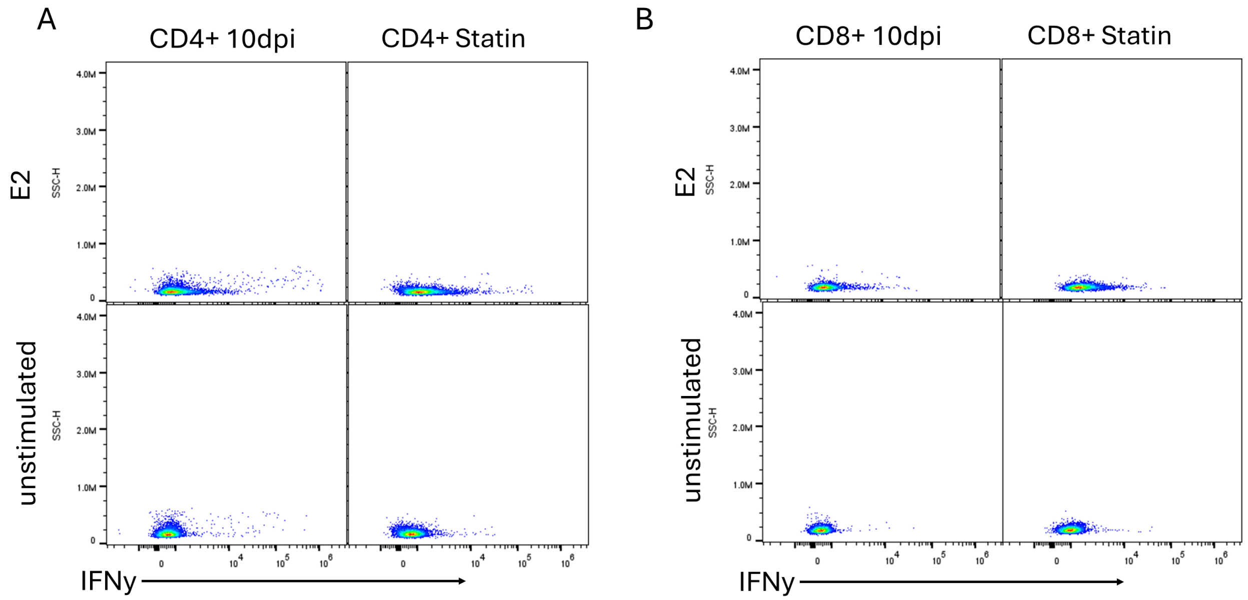


**Supplementary figure 8: Atorvastatin treatment modulates IFNy secretion in T cells following ONNV infection.** A and B show representative flow cytometry plots IFNy expression in CD4+ and CD8+ cells from lymph nodes of ONNV infected mice with or without statin treatment. These cells were stimulated ex vivo with ONNV-E2-Lfn or were left unstimulated as controls.

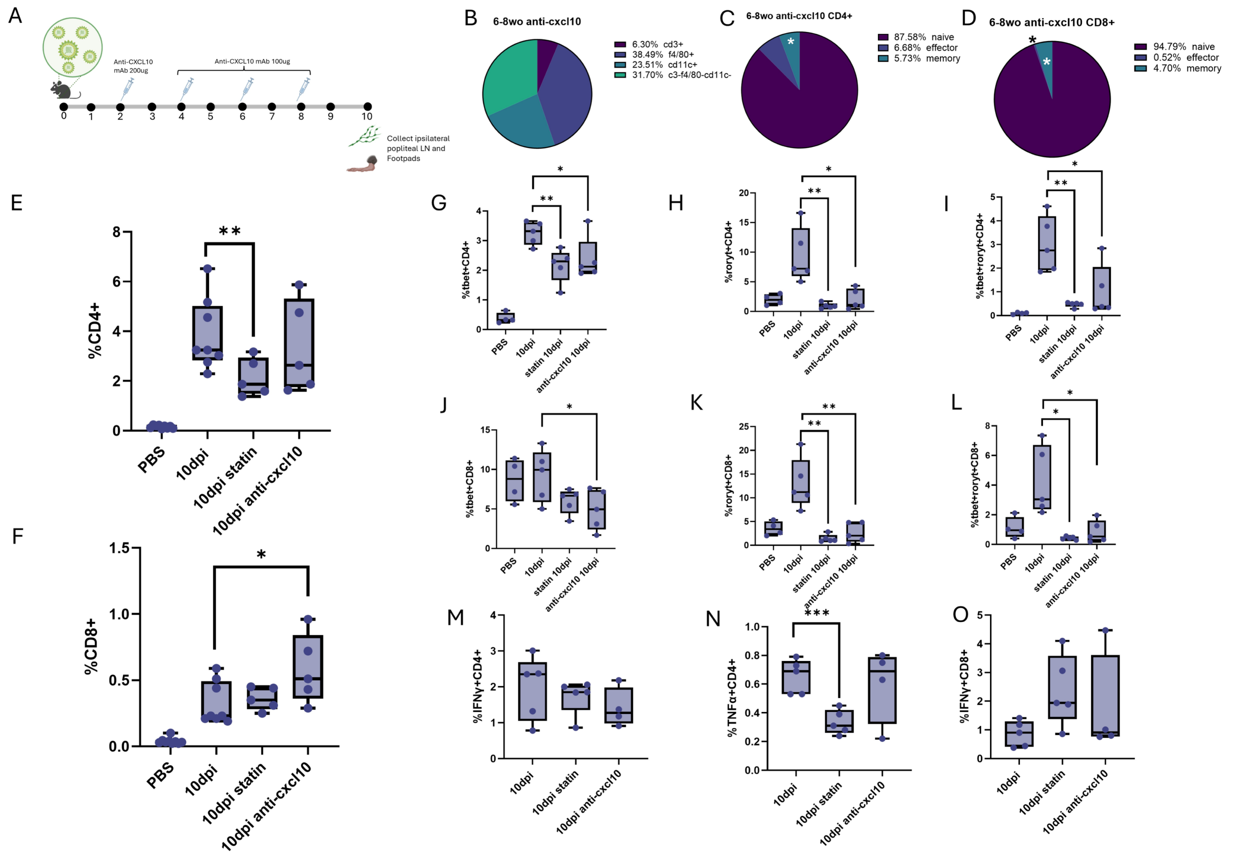


**Supplementary figure 9: Anti-CXCL10 antibody treatment reduces CD4+ T cell infiltration and attenuates proinflammatory T cell programming during ONNV infection.** 6-8wo mice were treated with IP doses of anticxcl10 antibody followed by collection of footpads and lymph nodes at 10dpi for flow cytometric analysis. A shows a schematic representation of the treatment strategy. B shows the average frequency (N=5) of CD3+, F480+ (Macrophages) and CD11c+ (Dendritic cells) in the footpads of statin treated mice. C and D show the average frequency (N=5) of naïve (CD44-CD62L+), effector (CD44+ CD62L-) and central memory (CD44+C62L+) CD4+ (C) and CD8+ (D) T cells. In each instance (E-F) cells from uninfected, infected without treatment, and statin treatment conditions are plotted alongside to enable comparison. E and F respectively show frequency of CD4+ and CD8+ cells in the footpad. G-I and J-L show frequency of Tbet+, Roryt+ or double positive CD4+ or CD8+ cells (N=5). M-O quantify IFNy+ CD4+, TNFa+CD4+ and IFNy+CD8+ cells in the lymph node after ex vivo stimulation with ONNV-E2-Lfn (N=5). Statistical significance was calculated using two-sample T-tests and the indicated significance is relative to relative to non-statin treated mice at 10dpi, where $*p < 0.05$, $**p < 0.01$, $***p < 0.001$, $****p < 0.0001$.

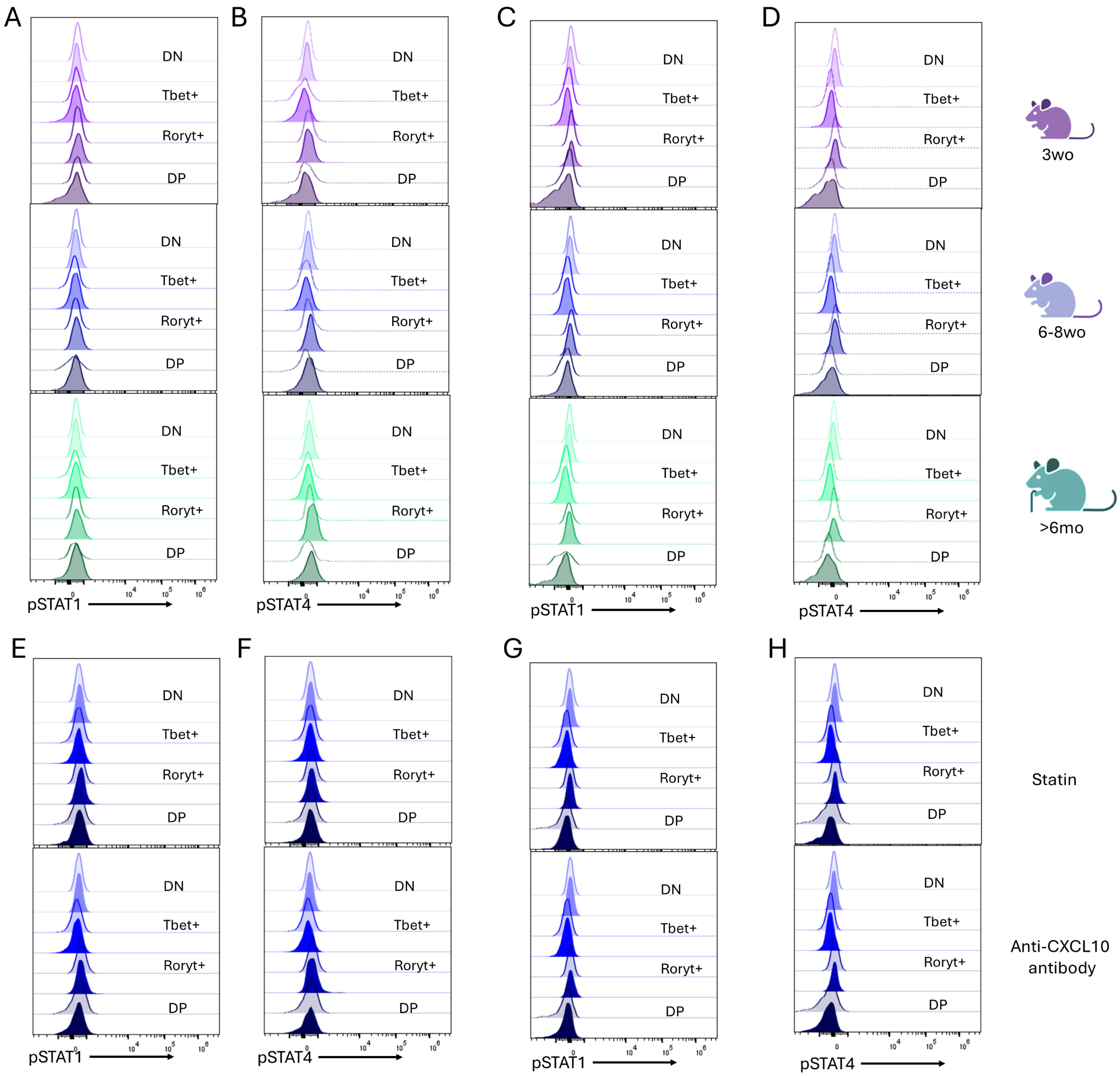


**Supplementary Figure 10: ONNV infection and blocking CXCL10 does not affect phosphorylation of STAT1 and STAT4.** 3wo, 6-8wo and >6mo mice were infected with ONNV and lymph nodes were collected at 10dpi for flow cytometric analysis. Histograms for the expression of pSTAT1 (A, C, E, G), pSTAT4 (B, D, F, H) were generated by combining expression in cells from N=3 mice. Cells gated on CD4+ (A and B) and CD8+ (C and D) expression and split into Tbet+, Roryt+, double positive (DP) or double negative (DN)

categories, and the dotted lines represent corresponding uninfected controls. E-H compares expression in ONNV infected mice (translucent colors) and either atorvastatin or Anti-cxcl10 antibody treated mice (opaque colors), as indicated, where E-F correspond to CD4+ and G-H correspond to CD8+ cells.

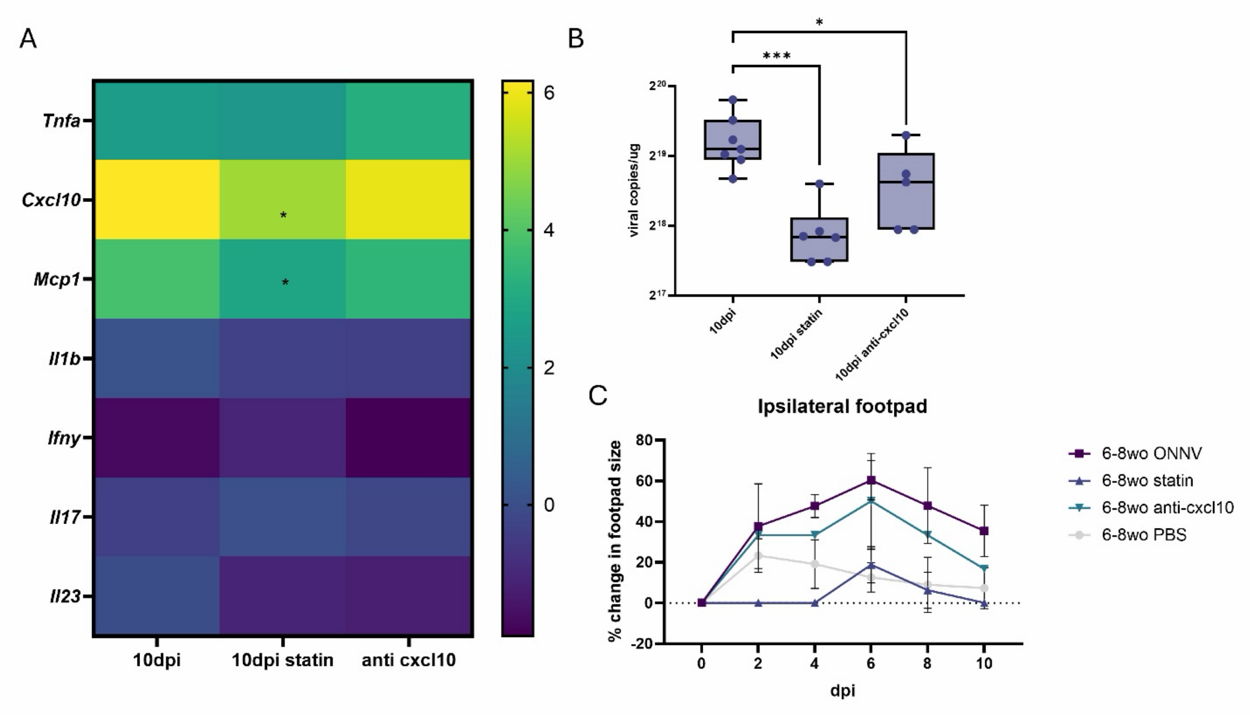


**Supplementary figure 11: Anti-CXCL10 antibody treatment reduces footpad swelling and viral burden at 10 dpi.** Following infection and anti-CXCL10 antibody treatment, 6-8wo mice footpads were subjected to qPCR. In each instance, infected without treatment, and statin treatment conditions are included alongside to enable comparison. A shows the average (across N=5-7) Log2-transformed fold change in expression of various genes relative to uninfected controls is shown. B shows footpad viral load measured by qPCR (N=5-7). C shows change in footpad size following ONNV infection and atorvastatin treatment (N=2-4). Statistical significance was calculated using two-sample T-tests and the indicated significance is relative to non-statin treated mice at 10dpi, where $*p < 0.05$, $**p < 0.01$, $***p < 0.001$, $****p < 0.0001$.
